# Local activation of CA1 pyramidal cells induces theta phase precession

**DOI:** 10.1101/2023.08.19.553985

**Authors:** Hadas E. Sloin, Lidor Spivak, Amir Levi, Roni Gattegno, Shirly Someck, Eran Stark

## Abstract

Hippocampal theta phase precession is involved in spatiotemporal coding and generating multineural spike sequences, but how precession originates remains unresolved. To determine whether precession can be generated directly in CA1 and disambiguate multiple competing mechanisms, we used optogenetic activation to impose artificial place fields in pyramidal cells of mice running on a linear track. More than a third of the CA1 artificial fields exhibited synthetic precession that persisted for a full cycle. In contrast, artificial fields in the parietal cortex did not exhibit synthetic precession. The findings are incompatible with precession models based on inheritance, spreading activation, dual-input, or inhibition-excitation summation. Thus, a precession generator resides locally within CA1.

## Introduction

Hippocampal theta phase precession is one of the most prominent examples of a temporal code. As an animal traverses a place field, spikes of pyramidal cells (PYRs) in the hippocampus and associated regions occur at progressively earlier theta phases (O’Keefe and Recce, 1993; Skaggs et al., 1996; Hafting et al., 2008). Precession also occurs outside the hippocampal formation (Jones and Wilson, 2005; Van Der Meer and Redish, 2011) and in relation to high frequency (Bush et al., 2022) and non-oscillatory (Eliav et al., 2018) bands. In the hippocampus, precession encodes space (Jensen and Lisman, 2000; Huxter et al., 2003; Maurer et al., 2014), time (Hirase et al., 1999; Pastalkova et al., 2008; Shimbo et al., 2021), objects (Robinson et al., 2017), and events (Terada et al., 2017). Precession is a potential mechanism for the generation of theta sequences (Skaggs et al., 1996; Chadwick et al., 2016; Reifenstein et al., 2021; George et al., 2023), which are involved in learning places and events (Dragoi and Buzsáki, 2006; Foster and Wilson, 2006; Drieu et al., 2018). Phase precession thus appears to be a general coding scheme and is a putative mechanism underlying learning (Jaramillo and Kempter, 2017).

Numerous models have been suggested for the generation of precession, which can be classified into five main categories (Jaramillo and Kempter, 2017; Drieu and Zugaro, 2019). “Inheritance” type models (Zugaro et al., 2005; Burgess et al., 2007; Hafting et al., 2008; Jaramillo et al., 2014) propose that precession is inherited from precessing neurons in the entorhinal cortex (EC) or CA3, which increase their excitatory drive earlier in each theta cycle (**Fig. 1A**). “Dual-input” models (Chance, 2012; Fernández-Ruiz et al., 2017) suggest that precession is generated by the gradual transition between early inputs from CA3 (at late theta phases) and later input from EC (at early theta phases; **Fig. 1B**). “Spreading activation” models (Jensen and Lisman, 1996; Tsodyks et al., 1996) suggest that precession results from feedforward excitation along a connected chain of place cells encoding the consecutive spatial regions that the animal passes through (**Fig. 1C**). Fourth, “inhibition-excitation summation” models (Kamondi et al., 1998; Magee, 2001; Harris et al., 2002; Mehta et al., 2002) generate precession by the summation of inhibitory theta currents and spatially-dependent depolarization, causing excitation to exceed inhibition earlier in subsequent theta cycles (**Fig. 1D**). Finally, in “dual-oscillator” models (O’Keefe and Recce, 1993; Bose et al., 2000; Lengyel et al., 2003), precession emerges from the superposition of theta and another faster rhythm which is triggered upon entering the place field (**Fig. 1E**). Faster oscillations could arise from a single-cell mechanism (e.g., within a PYR) or from local-circuit interactions (Bose et al., 2000; Chadwick et al., 2016) between PYRs and inhibitory interneurons (INTs). Despite the numerous models, contradictory experimental evidence hinders a clear understanding of how precession is generated (Zugaro et al., 2005; Harvey et al., 2009; Ravassard et al., 2013).

**Figure 1.**
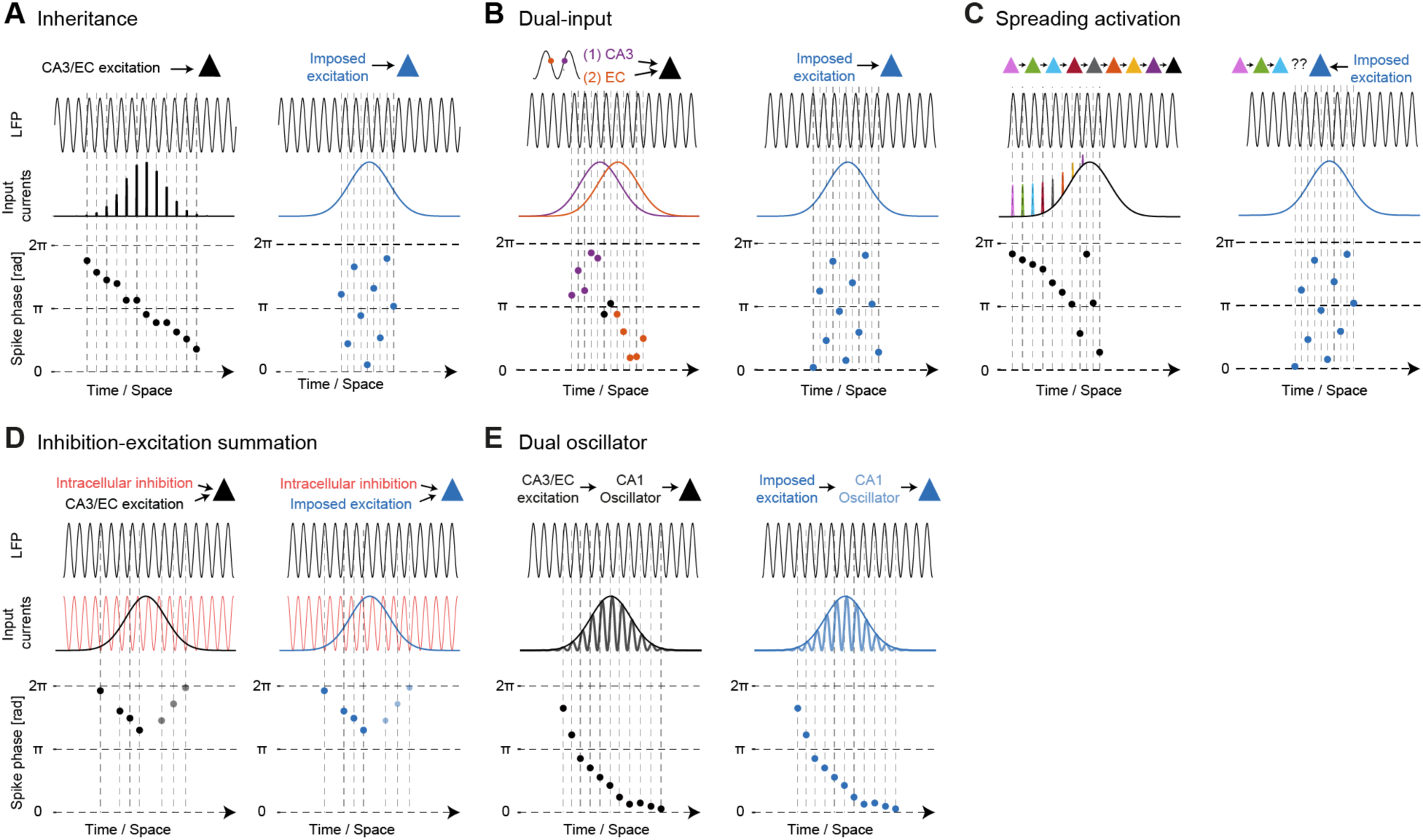
Artificial place fields can distinguish between models for the generation of theta phase precession. Every panel shows precession in spontaneous fields (left), and predictions for artificial fields (right) according to a different model. For simplicity, constant running speed is assumed. (**A**) CA1 PYRs inherit precession from precessing CA3/EC inputs. Imposed depolarization lacks theta-phase dependence, predicting no precession. (**B**) During place field traversal, excitation is first received from CA3 (at later theta phases) and then from EC (at early theta phases). Imposed depolarization from a single phase-independent source does not generate precession. (**C**) Precession is generated by spreading activation from a chain of connected place cells upstream in the trajectory of the animal. Imposing an artificial field in a cell not part of a matching chain does not induce precession. (**D**) Spatially dependent excitation surpasses ongoing inhibitory theta currents in early LFP theta cycles, creating half-cycle precession. In the second half of the field, spikes shift in the opposite direction (grey) or disappear due to adaptation. An artificially imposed field exhibits a similar pattern. (**E**) Excitatory inputs trigger a fast oscillator within CA1, which peaks earlier in each theta cycle. An imposed depolarization that triggers the fast oscillator produces precession.

## Results

### Pyramidal cell activation induces synthetic theta phase precession

We tested the validity of the various models for the generation of precession by inducing artificial place fields directly in CA1 PYR using focal optogenetic activation. In an artificial field, a depolarizing current is imposed upon PYRs, independent of the theta phase. The different models yield distinct predictions regarding the theta phase of the induced spikes. Assuming that the activated PYR carries no subthreshold or suprathreshold spatial activity, the predictions are as follows. If precession is inherited to CA1 PYR from precessing CA3/EC inputs, artificial place fields would not precess, because the imposed depolarization does not precess (**Fig. 1A**, right). If precession is created by dual-input arriving first from CA3 and then from EC, artificial place fields would not precess since the imposed excitation is indifferent to phase (**Fig. 1B**). Likewise, imposed activation of a cell without exciting the specific chain of presynaptic neurons matching the trajectory of the animal will not induce spreading activation, and thus will not generate precession (**Fig. 1C**). In contrast, inhibition-excitation summation models predict that imposed excitation would result in precession during the first half of the place field due to the concurrent presence of inhibitory theta inputs (**Fig. 1D**). In the second half-field, precession would either disappear due to adaptation (Harris et al., 2002) or shift in the opposite direction (Yamaguchi et al., 2002; Wang et al., 2020). Finally, if the imposed excitation triggers a faster-than-theta oscillator, dual-oscillator models predict that the induced spikes could undergo a complete precession cycle (**Fig. 1E**).

To create a place-field like imposed excitation in CA1 PYRs, we employed focal closed-loop optogenetic activation in freely-moving mice (**Fig. S1A**). As CaMKII::ChR2 mice (n=5; **Table S1**) ran on a 150 cm linear track, illumination was provided based on real-time orientation and position during half of the same-direction trials (**Fig. 2A**). An example PYR rarely fired during Control trials but exhibited a position-dependent firing rate increase during Light trials (**Fig. 2B**). Among active and stable CA1 PYRs, 676/1,095 (62%) showed firing rate increases that matched the illumination pattern (**Fig. 2C**; **Table S2**). We categorized PYRs based on firing rate gain and correlation with the illumination pattern (p<0.05, permutation test; **Fig. 2D**; **Fig. S1F**). 286/1,095 (26%) PYRs exhibited artificial place fields which were present only during Light but not during Control trials (p<1.11×10^-16^, binomial test). We quantified spiking within the illuminated region by field selectivity, the firing rate during illumination divided by the firing rate out of the light limits. Selectivity of artificial fields was above one specifically during Light trials (p=2.6×10^-48^, Wilcoxon’s test; **Fig. 2E**). Artificial fields appeared only locally, in PYRs recorded by the illuminated shank (p<1.11×10^-16^; **Fig. S1G**). Hence, closed-loop illumination of CA1 PYRs imposes local artificial place fields.

**Figure 2.**
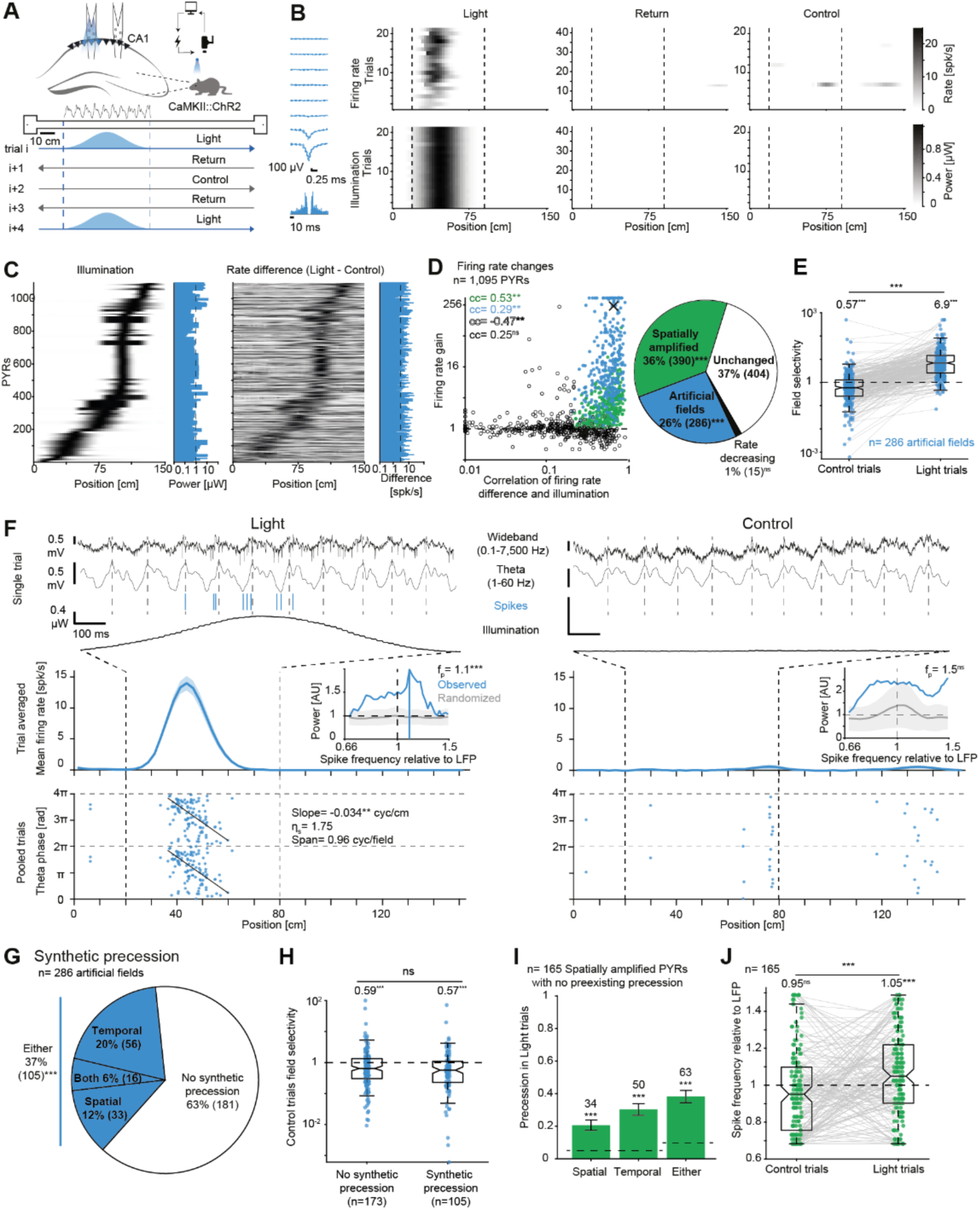
Pyramidal cell activation induces synthetic theta phase precession. (**A**) Experimental design. (**B**) Focal CA1 illumination generates an artificial place field in a PYR during left-to-right runs. Dashed lines, light limits. Left, Wideband spike waveform (mean±SD) and autocorrelation. (**C**) Position-dependent illumination increases PYR firing. Histograms, peak power/rate difference. (**D**) Illumination generates artificial fields and amplifies preexisting fields. cc, rank correlation coefficient. ns/**: p>0.05/p<0.01, permutation test. X, example PYR shown in **BF**. Right, Illumination induces artificial fields in a quarter of PYRs. Here and in **GI**, ns/***: p>0.05/p<0.001, binomial test. (**E**) Field selectivity is above one during Light but not Control trials. Here and in **HJ**, ***: p>0.05/p<0.001, Wilcoxon’s test. (**F**) Focal activation induces synthetic precession. Top, Example Control and Light trials. Middle, Mean±SEM firing rate. Bottom, Theta phase vs. animal position at each spike. **: p<0.01, permutation test. Inset, Spectrum of the observed spike train and of phase-randomized spikes. ns/***: p>0.05/p<0.001, constrained randomization test. (**G**) Illumination induces precession in one-third of the artificial fields. (**H**) Field selectivity in artificial fields. (**I**) Over a third of spatially amplified PYRs precess during Light trials. Horizontal lines, chance levels; error bars, SEM. (**J**) Spatially amplified PYRs exhibit higher spike frequency during Light trials.

The induced spikes in an example artificial field occurred earlier in consecutive theta cycles, indicating synthetic precession (**Fig. 2F**). Among artificial place fields, 105/286 (37%) PYRs exhibited synthetic precession (p<1.11×10^-16^; binomial test; **Fig. 2G**), including 49 PYRs with synthetic spatial precession (p=7.68×10^-5^) and 72 PYRs with synthetic temporal precession (p<4.54×10^-14^; **Fig. S2**; **Fig. S3A**). In 75/286 (26%) of the artificial fields, the induced spikes were locked to a specific theta phase (p<1.11×10^-16^; binomial test; **Fig. S3BC**). Synthetic precession was faster than spontaneous precession (p<0.046, U-test; **Fig. S3D**). However, precession effect size did not differ between spontaneous and synthetic precession (p>0.11, U-test; **Fig. S3E**). Baseline activity across the track but not depth within CA1 str. pyramidale differed between PYRs with and without synthetic precession (**Fig. S4AB**; **Table S3**). Thus, locally imposed artificial place fields exhibit synthetic precession.

Depolarization may unmask place fields in regions with preexisting subthreshold excitatory drive (Lee et al., 2012; McKenzie et al., 2021; Valero et al., 2022). In contrast to the previous studies that applied excitation in contiguous blocks of trials, the artificial fields generated in the alternating Light trials (**Fig. 2A**) did not exhibit spatial selectivity to the illuminated region during Control trials (**Fig. 2E**). Furthermore, the firing rate within the light limits during Control trials was 0.26 spk/s for PYRs with artificial place fields, lower than for PYRs showing no change in firing rate (3.2 spk/s; **Fig S5**). In contrast, field selectivity of artificial fields with synthetic precession was not higher than fields without synthetic precession (p=0.33, U-test; **Fig. 2H**). Thus, artificial place fields do not unmask subthreshold fields, and the likelihood of synthetic precession is not related to preexisting spatial selectivity.

In addition to the artificial place fields, 390/1,095 (36%) PYRs were “spatially amplified”, i.e., were active during Control trials and increased spiking during Light trials (**Fig. 2D**). Of the 165 spatially amplified PYRs which did not exhibit precession during Control trials, 63/165 (38%) PYRs exhibited precession during Light trials (p<1.11×10^-16^, binomial test; **Fig. 2I**). Furthermore, spatially amplified PYRs exhibited increased spike frequency relative to LFP, indicating temporal precession (p=2×10^-4^, Wilcoxon’s test; **Fig. 2J**). Thus, focal activation of CA1 PYRs induces synthetic precession in both artificial and preexisting place fields. Considering the model predictions, synthetic precession cannot be generated by the dual-input (**Fig. 1B**) or spreading activation (**Fig. 1C**) models.

### Synthetic precession spans a complete theta cycle and does not slow down or reverse direction

Both inhibition-excitation summation and dual-oscillator models predict synthetic precession, but the specifics differ. Inhibition-excitation summation models predict synthetic precession spanning half a theta cycle (**Fig. 1D**), whereas dual-oscillator models predict precession could span a full theta cycle (**Fig. 1E**). In example spontaneous and artificial fields, spontaneous precession spanned approximately half a theta cycle, while synthetic precession spanned a full cycle (**Fig. 3A**). In the cohort of artificial fields, synthetic precession spanned a median [interquartile interval, IQR] of 0.96 [0.6 1.43] cycles, not different from a full theta cycle (p=0.25, Wilcoxon’s test; **Fig. 3B**). Spontaneous precession spanned 0.78 [0.63 1.12] cycles, more than half a cycle (p=2.4×10^-47^, Wilcoxon’s test). There was no consistent difference between the spans of synthetic and spontaneous precession (p=0.088, U-test; **Fig. 3B**).

**Figure 3.**
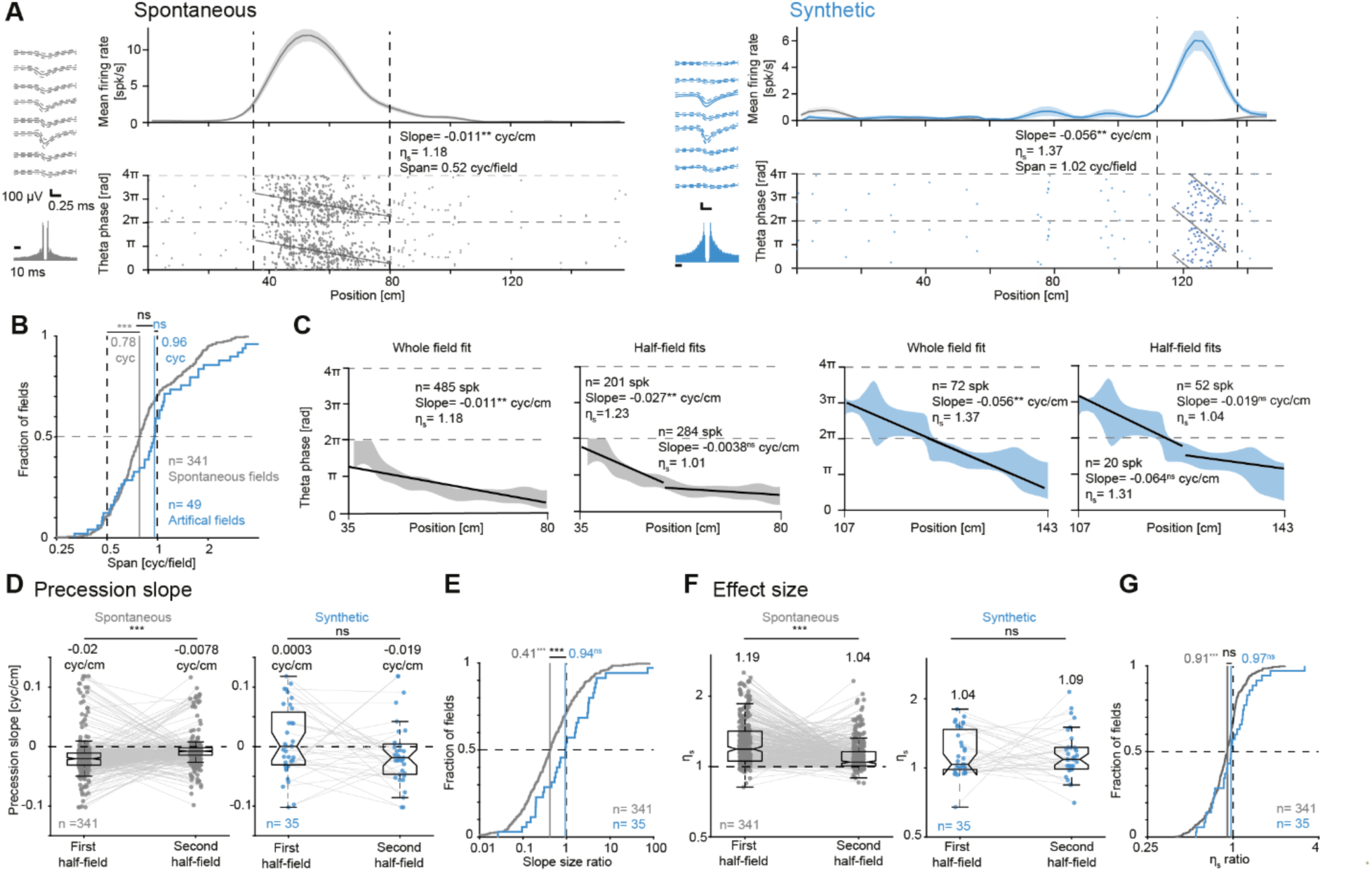
Synthetic precession spans a complete theta cycle and does not slow down or reverse direction. (**A**) Left, Example CA1 PYR with spontaneous precession. Right, Example PYR with synthetic precession. All conventions are the same as in Fig. 2F. (**B**) Synthetic precession span is not different from a full cycle. Spontaneous precession spans more than half a cycle. Lined ns/***: p>0.05/p<0.001, Wilcoxon’s test comparing to a null of 1 and 0.5, for synthetic and spontaneous precession. (**C**) In the example PYRs, the first half-field exhibits higher slope and effect size during spontaneous but not during synthetic precession. Gray/blue patches, mean±SD theta phase of spikes as a function of position. Black lines, circular-linear fits. (**D**) Precession slope in the two halves of spontaneous and synthetic fields. Here and in **F**, lined ns/***: p>0.05/p<0.001, Wilcoxon’s test. (**E**) Precession slope in the second half-field, divided by the first. Values smaller than one indicate that precession slows down. ns/***: p>0.05/p<0.001, Wilcoxon’s test compared to a unity null. (**F**) Effect size is smaller in the second half-field of spontaneous but not of synthetic precession. (**G**) Spatial effect size ratio, the effect size for the second half-field divided by the effect size for the first half-field.

Previous studies with rats showed that spontaneous precession changes direction (Yamaguchi et al., 2002; Wang et al., 2020) in the second half of the place field, consistent with predictions of some versions of the excitation-inhibition summation model (**Fig. 1D**, light dots). We found that spontaneous precession slowed down and diminished in the second half of the place field, but synthetic precession did not (**Fig. 3C**). Overall, spontaneous precession slopes were more negative in the first half-field compared with the second (p=5.4×10^-11^, Wilcoxon’s test; **Fig. 3D**, left). In contrast, synthetic precession slopes did not slow down, and were −0.019 [−0.047 0.0044] cyc/cm in the second half-field (**Fig. 3D**, right). Consequently, the ratio between the slope sizes in the second and first half-fields was smaller in spontaneous compared with synthetic precession (p=0.0046, U-test; **Fig. 3E**). The difference between slope size ratios illustrates that spontaneous precession slows down, while synthetic precession does not.

A prediction of other versions of the excitation-inhibition summation model (**Fig. 1D**) is that precession will diminish in the second half-field (Harris et al., 2002; Mehta et al., 2002; Yamaguchi et al., 2002). While spontaneous fields indeed displayed smaller precession effect size in the second half-field (p=2.8×10^-15^, Wilcoxon’s test), the effect size did not decrease in the second half of artificial fields (p=0.6, Wilcoxon’s test; **Fig. 3F**). Accordingly, the ratio between precession effect sizes in the second and first half-fields was distinct from one among spontaneous (p=1.7×10^-10^, Wilcoxon’s test) but not synthetic fields (p=0.5; **Fig. 3G**). Thus, precession slope and strength do not diminish in the second half of the artificial field, indicating that synthetic precession cannot be generated by an inhibition-excitation summation model (**Fig. 1D**).

### Focal pyramidal cell activation slows preexisting spontaneous precession

While artificial fields are not unmasked subthreshold fields (**Fig. 2E**; **Fig. S5**), the synthetic precession observed during local activation of PYR could be generated locally or inherited from an upstream source (**Fig. 1A**). To examine whether the generator of spontaneous precession (**Fig. 4A**, left) resides in CA1, we investigated the effects of local activation on place cells with preexisting precession within the illuminated region. A priori, if precession is inherited (e.g., from CA3/EC), local CA1 activation may reduce precession prevalence and effect size but would not modify precession speed, as the precession generator itself is not affected (**Fig. 4A**, middle). In contrast, if precession originates in CA1, local activation may modify precession speed by directly biasing the generator (**Fig. 4A**, right).

**Figure 4.**
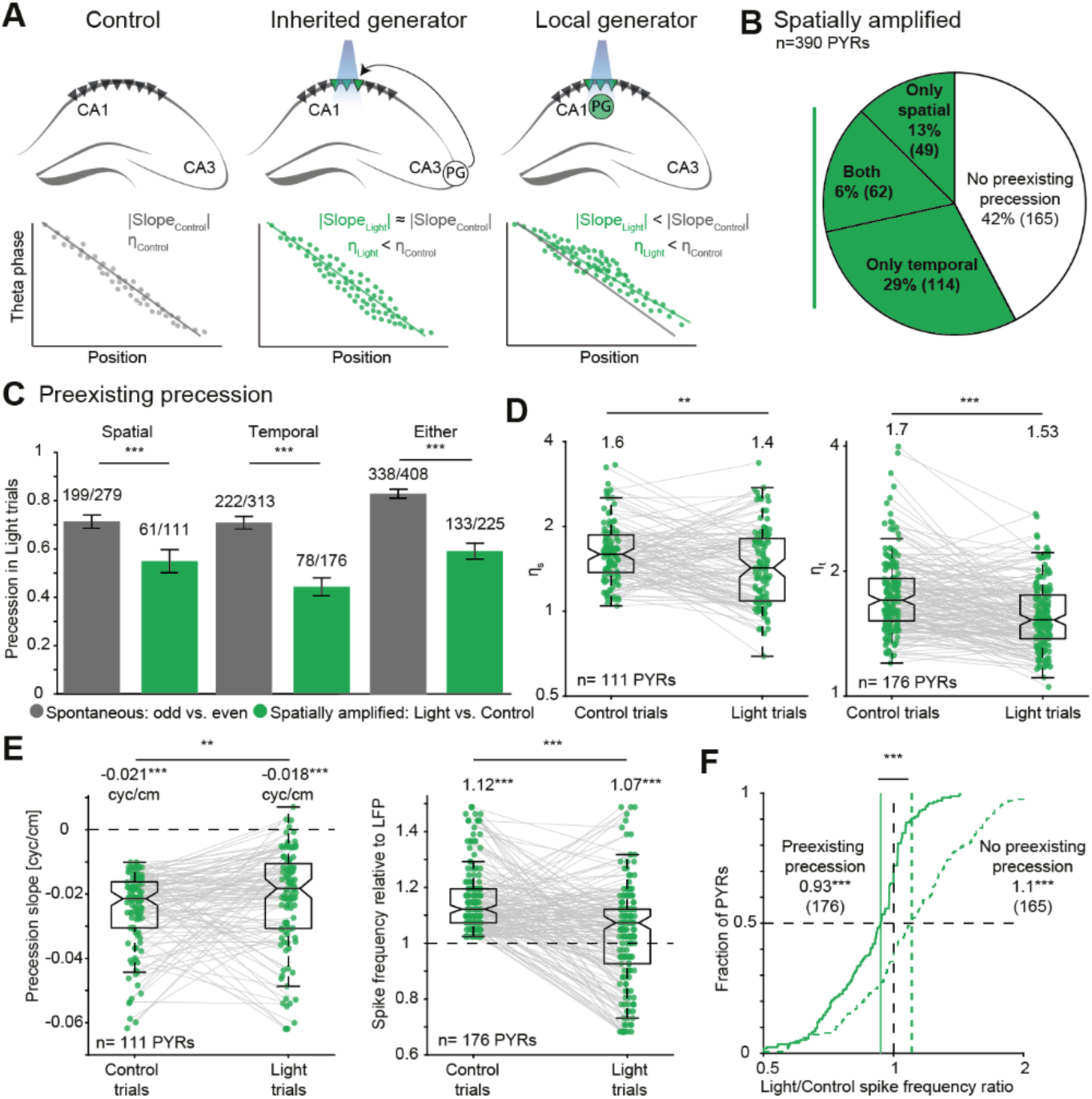
Focal pyramidal cell activation slows preexisting spontaneous precession. (**A**) Possible outcomes of activating PYRs with preexisting precession, based on inheritance (e.g., from CA3) as opposed to local precession generation (PG). (**B**) Spontaneously active PYRs that exhibit a light-dependent rate increase include units with and without preexisting precession. (**C**) Illumination reduces precession prevalence among PYRs with preexisting precession. Spatially amplified Light vs. Control, precession during Light/Control for spatially amplified PYR with preexisting precession. Spontaneous odd vs. even, precession during odd/even trials within spontaneous place fields. Lined ***: p<0.001, G-test. (**D**) Focal activation reduces the effect size of preexisting precession. Here and in **E**, lined **/***: p<0.01/p<0.001, Wilcoxon’s test. (**E**) Activation slows preexisting precession during Light compared with Control trials. Left, Precession slope size is smaller. ***, p<0.001, Wilcoxon’s test compared with a zero null (one-tailed). Right, Spike frequency relative to LFP is smaller. Here and in **F**, ***: p<0.001, Wilcoxon’s test compared to unity null (one-tailed). (**F**) Activation exerts opposite effects on spike frequency of precessing (continued line) and non-precessing spatially amplified PYRs (dashed line). Lined ***: p<0.001, U-test.

We found that 225/390 (58%) of the spatially amplified PYRs exhibited preexisting precession within light limits (**Fig. 4B**). The prevalence of precession was reduced during Light compared with Control trials (133/225; 59%; **Fig. 4C**, green), beyond the reduction observed in spontaneous place fields during no-light odd-vs. even-numbered trials (338/408; 83%; p<1.1×10^-16^, G-test; **Fig. 4C**, grey). Furthermore, precession effect sizes were lower during Light compared with Control trials (p<9.1×10^-3^; Wilcoxon’s test; **Fig. 4D**). Activation reduced spatial precession slope (p=0.0084, Wilcoxon’s test) and spike frequency relative to the LFP (p=2.2×10^-10^, Wilcoxon’s test; **Fig. 4E**).

The findings (**Fig. 4DE**) contrast with the increased spike frequency among other spatially amplified PYRs (**Fig. 2J**). Specifically, activation increased spike frequency in PYRs without preexisting precession (p=0.018, Wilcoxon’s test) but decreased frequency in PYRs with preexisting precession (p=3.9×10^-7^), exerting opposite effects (p*=*6.9×10^-12^, U-test; **Fig. 4F**). The impact of activation on precession speed is at odds with inheritance models (**Fig. 1A**; **Fig. 4A**, center), suggesting that the generator underlying spontaneous precession resides in CA1.

### Focal PYR activation induces a rate increase without precession in CA1 INTs and parietal cortex PYRs

Various factors within CA1 may contribute to precession generation, including local-circuit PYR-INT interactions (Bose et al., 2000; Chadwick et al., 2016). Consistent with the circuit models, CA1 INTs may exhibit spontaneous precession (Maurer et al., 2006; Ego-Stengel and Wilson, 2007), and silencing INTs affects spontaneous precession (Royer et al., 2012; Grienberger et al., 2017). We therefore examined whether focal PYR activation induces synthetic precession in local-circuit INTs. 85/158 (54%) of the local INTs, recorded on the illuminated shank, exhibited position-dependent firing rate increase (**Fig. 5AB**; **Fig. S6AB**; **Table S4**). Only 3/158 (2%) INTs exhibited artificial fields (p=0.99, binomial test; **Fig. 5C**; **Fig. S6CD**), precluding the characterization of synthetic precession in INTs. A total of 68 local INTs were spatially amplified (**Fig. S6E**) and did not exhibit preexisting precession (**Fig. 5D**), of which only 5 INTs (7%) exhibited precession during Light trials (p=0.8, binomial test; **Fig. 5E**). Spike frequency relative to LFP of CA1 INT did not change between Control and Light trials (p=0.78, Wilcoxon’s test; **Fig. 5F**). Thus, while a position-dependent rate increase is transferred from PYRs to INTs, synthetic precession does not necessarily involve precession in local INT as suggested by some local circuit models (Bose et al., 2000; Chadwick et al., 2016). Notably, circuit models may generate synthetic precession if e.g., non-local INTs are involved.

**Figure 5.**
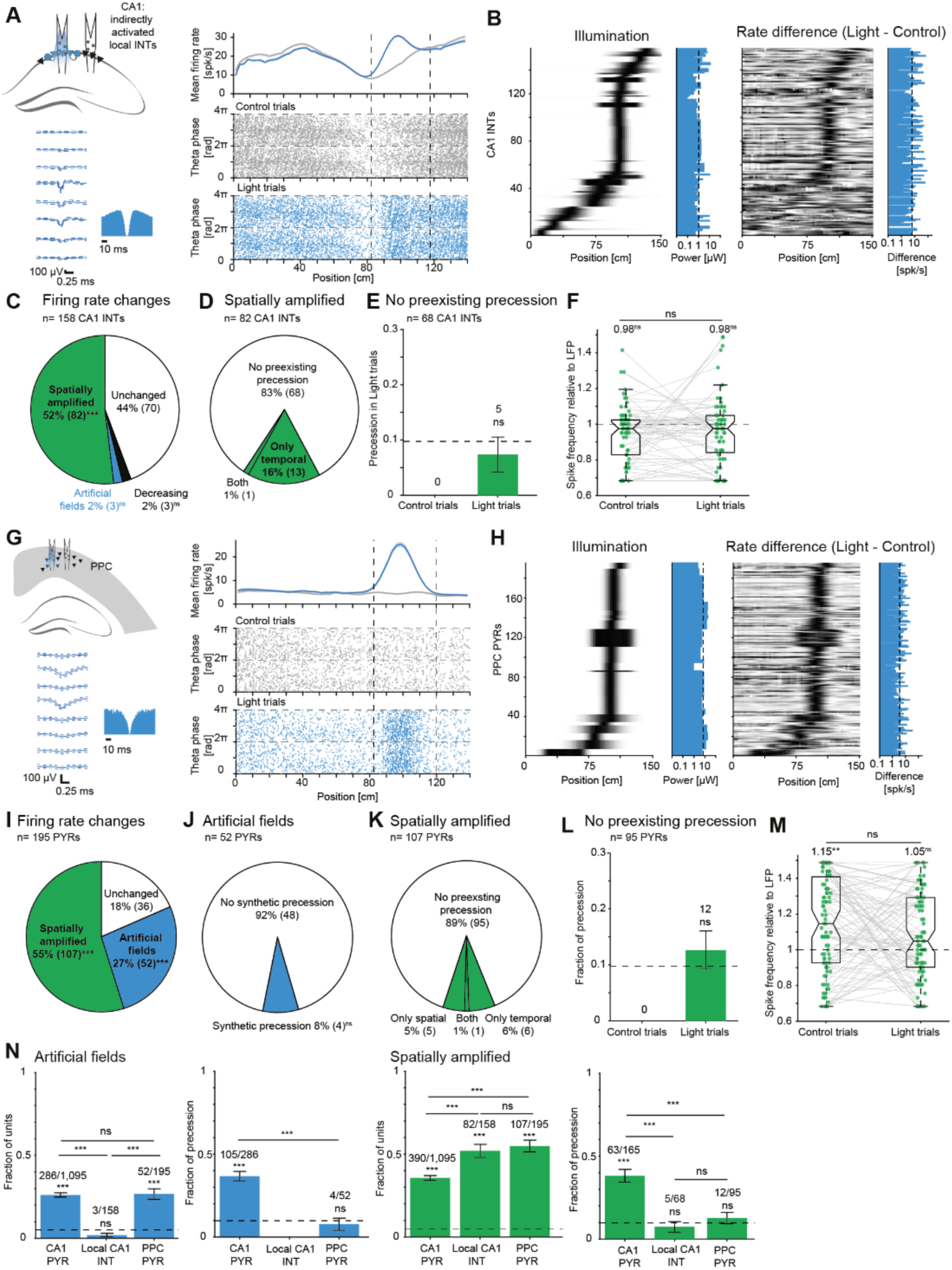
Focal PYR activation induces a rate increase without precession in CA1 INTs and parietal cortex PYRs. (**A**) Position-dependent CA1 PYR illumination indirectly activates local INTs. (**B**) PYR activation increases firing rates of INT on the illuminated shank. (**C**) PYR activation does not generate artificial fields in local INTs. (**D**) Precession prevalence during Control trials in spatially amplified INTs. (**E**) Spatially amplified INTs do not precess during Light trials. Here and in **L**, ns: p>0.0975, binomial test. (**F**) Local INT spike frequency relative to LFP is not higher during Light compared with Control trials. (**G**) In PPC of CaMKII::ChR2 mice, focal closed-loop illumination activates PPC PYRs. (**H**) Illumination activates local PYRs. (**I**) Activation induces artificial fields in PPC PYRs. (**J**) PPC PYRs with artificial fields do not exhibit synthetic precession. (**K**) Precession prevalence during Control trials in spatially amplified PPC PYRs. (**L**) PPC PYR activation does not affect precession prevalence in spatially amplified PYRs. (**M**) PPC PYR spike frequency relative to LFP is not increased during Light trials. (**N**) Synthetic precession occurs in light-activated CA1 PYRs, but not in indirectly-activated CA1 INTs or in light-activated PPC PYRs. ns/***: p>0.05/p<0.001, binomial test. Lined ns/*/**/***: p>0.05/p<0.05/p<0.01/p<0.001, G-test.

We found that in CA1, an imposed excitation of PYR is sufficient to generate precession. To investigate whether a rate increase combined with theta may suffice for synthetic precession, we repeated the imposed excitation experiment in the posterior parietal cortex (PPC; n=22 sessions in two CaMKII::ChR2 mice; **Fig. 5G**; **Fig. S7AB**; **Table S5**). Hippocampal theta oscillations are volume-conducted to the PPC, where neurons are phase-locked to specific theta phases (Sirota et al., 2008). PPC neurons exhibit spatially-modulated activity (Krumin et al., 2018), but no spontaneous phase precession (**Fig. S8**).

Focal PYR activation in the PPC induced position-dependent firing rate increases (**Fig. 5H**), with 52/195 (27%) PPC PYRs showing artificial place fields (p<1.11×10^-16^, binomial test; **Fig. 5I**). However, synthetic precession was observed at chance level (4/52 PPC artificial fields, 8%; p=0.76, binomial test; **Fig. 5J**). Among spatially amplified PPC PYRs (**Fig. 5K**), precession occurred in 12/95 (13%) PYRs, not above chance (p=0.21, binomial test; **Fig. 5L**). Furthermore, spike frequency relative to LFP of spatially amplified PPC PYR did not increase between Control and Light trials (p=0.11, Wilcoxon’s test; **Fig. 5M**). To summarize, PPC activation generates artificial place fields but is insufficient to create synthetic precession. Together (**Fig. 5N**), the findings suggest that synthetic precession requires synaptic theta or depends on cellular-network properties found in CA1 but not in the PPC.

## Discussion

We found that imposing artificial place fields on CA1 PYRs generates synthetic precession, which is incompatible with spreading activation and dual-input models. Synthetic precession does not diminish or reverse in the second half of the field, inconsistent with inhibition-excitation summation models. Local activation slows preexisting precession, suggesting that the generator of spontaneous precession is not inherited but rather resides in CA1. The synthetic precession mechanism does not involve precession of local-circuit CA1 INTs but does require properties absent in the PPC, e.g., synaptic arrival of theta, the circuit architecture of CA1, or biophysics unique to CA1 PYRs.

Among the models proposed for precession generation, the results are consistent with dual-oscillator models (O’Keefe and Recce, 1993; Bose et al., 2000; Lengyel et al., 2003; Chadwick et al., 2016), which require theta and faster oscillations. PYR membrane potentials may demonstrate faster-than-theta oscillations due to interacting amplifying and resonant currents (Hutcheon and Yarom, 2000; Izhikevich, 2006). Alternatively, a PYR may exhibit fast oscillations arising from local circuit interactions (Bose et al., 2000; Stark et al., 2013; Chadwick et al., 2016). Synthetic precession may originate from a yet unidentified mechanism that meets the following constraints: rate dependence, completion of a full cycle, independence of local INT precession, and CA1 specificity. The existence of synthetic precession indicates that in the face of ongoing theta, local excitation of CA1 circuity is sufficient for generating precession, transforming a rate code into a temporal code.

Synthetic precession carries implications for the functional role of precession in generating theta sequences and learning (Jaramillo and Kempter, 2017; Drieu and Zugaro, 2019). While precession has been proposed to generate theta sequences (Skaggs et al., 1996; Chadwick et al., 2016; Reifenstein et al., 2021; George et al., 2023), others suggested that both precession and sequences arise from preconfigured connectivity of PYRs (Jensen and Lisman, 1996; Mizuseki and Buzsáki, 2013; Venditto et al., 2019). We found that precession can be generated by activating arbitrary CA1 PYRs at random positions along the track. Since the activated PYRs are not expected to have prior synaptic connections with one another, preconfigured connectivity is unlikely to suffice for generating synthetic precession. Synthetic precession may therefore generate theta sequences between previously unassociated PYRs, potentially facilitating the learning of novel sequences (Wang et al., 2015; George et al., 2023).

## Acknowledgments

We thank György Buzsáki, Kamran Diba, Amit Marmelshtein, Shaked Palgi, Horacio G. Rotstein, and Michaël Zugaro for constructive comments. This work was supported by the United States-Israel Binational Science Foundation (BSF; 2015577); by the European Research Council (679253); by the Israel Science Foundation (638/16); by the Canadian Institutes of Health Research (CIHR), the International Development Research Centre (IDRC), the Israel Science Foundation (ISF) and the Azrieli Foundation Grant #2558/18; and by the Rosetrees Trust (A1576).

## Author contributions

E.S. conceived the study. H.E.S. and E.S. constructed optoelectronic probes and implanted animals. H.E.S. carried out the experiments. H.E.S., L.S., A.L., R.G., S.S., and E.S. analyzed data. H.E.S. and E.S. wrote the manuscript with input from all authors.

## Materials and Methods

### Experimental animals

Five freely-moving mice, four males and one female, were used in this study (**Table S1**). All mice expressed ChR2 under the CaMKII promoter. Three mice were double transgenics, offspring of CaMKII-Cre driver males (JAX #005359, The Jackson Labs) and ChR2 reporter females (Ai32; JAX #012569). Two mice were double transgenic and hybrid (Sloin et al., 2022a), obtained by crossing an FVB/NJ female (JAX # 001800) with a CaMKII::ChR2 male. All animals were healthy and not used for other procedures. At the time of implantation, animals aged 10-30 weeks and weighed 24.2-33.7 g. After implantation, mice were single housed to prevent damage to the implanted apparatus and held on a reverse dark/light cycle (dark phase from 8 AM until 8 PM). All animal handling procedures were in accordance with Directive 2010/63/EU of the European Parliament, complied with Israeli Animal Welfare Law (1994), and were approved by Tel Aviv University Institutional Animal Care and Use Committee (IACUC #01-16-051 and #01-21-018).

### Probes and surgery

Every mouse was implanted with a multi-shank silicon probe attached to a movable microdrive and equipped with optical fibers following previously described procedures (Stark et al., 2012; Noked et al., 2021). The probes used were Stark64 (Diagnostic Biochips; one mouse), Buzaski32 (NeuroNexus; one mouse), and Dual-sided64 (Diagnostic Biochips; three mice). The Stark64 probe consists of six shanks, spaced horizontally 200 µm apart, with each shank consisting of 10-11 recording sites, spaced vertically 15 µm apart. The Buzaski32 probe consists of four shanks, spaced horizontally 200 µm apart, with each shank consisting of eight recording sites, spaced vertically 20 µm apart. The Dual-sided64 probe consists of two dual-sided shanks, spaced horizontally 250 µm apart, with each shank consisting of 16 channels on each side (front and back), spaced vertically 20 µm apart.

Probes were implanted in the posterior parietal cortex (PPC) above the right hippocampus (PA/LM, - 1.6/1.1 mm; 45° to the midline, 0° to the vertical) under isoflurane (1%) anesthesia as previously described (Stark et al., 2012).

### Behavior and recording sessions

Full recovery from surgery was defined when weight ceased to decrease and the head was held autonomously. Following recovery, animals were placed on a water-restriction schedule that guaranteed at least 40 ml/kg of water (corresponding to 1 ml per 25 g mouse) on every recording day. During linear track running, data were recorded from the PPC of two mice and from hippocampal region CA1 of four mice; one mouse was used in both regions (**Table S1**). All hippocampal sessions were from the CA1 pyramidal cell layer, recognized by the appearance of multiple high-amplitude units and iso-potential spontaneous ripple events. Recordings were carried out five days a week, and animals received free water on the sixth day. After one to five recording sessions, the probe was translated vertically downwards by up to 70 µm.

Neuronal activity was recorded in 4.5 [1.9 10.1] hour sessions (median [interquartile range, IQR]). At the beginning of every session, baseline neural activity was recorded while the animal was in the home cage for 15-90 min. The animal was then placed on a 150 cm linear track that extended between two 10 x 10 cm square platforms. Each platform included a water delivery port. The water-restricted mice were trained to repeatedly traverse the track for a water reward of 3-10 µL. Mice ran 169 [133 200] one-direction trials during about one hour (**Table S2**, **Table S5**). Animals were equipped with a 3-axis accelerometer (ADXL-335, Analog Devices) for monitoring head movements. Head position and orientation were tracked using head-mounted LEDs of two different colors (green and red), captured by a machine vision camera (ace 1300-1200uc, Basler). Animal kinematics was estimated in real-time by a rigid-body Kalman filter algorithm running on the data acquisition machine by the “Spotter” program (Gaspar et al., 2019).

### Closed-loop illumination based on animal kinematics

The system was engineered to allow closed-loop illumination based on kinematic parameters of a freely-moving subject, position and head orientation (**Fig. S1A**). The hardware component of Spotter produced analog voltage signals proportional to the kinematics, which were routed to a digital signal processor (DSP; RX8, Tucker-Davis Technologies). Before every session, a kinematic illumination model was generated in MATLAB and uploaded to the DSP, coding voltages that are proportional to the LED power at every position and head orientation. At every instant, the DSP compared the real-time position and orientation to the model and issued a corresponding voltage command to a linear current source (Stark et al., 2012). The current source converted the voltages to currents that passed via an ultra-light Litz cable to a head-mounted LED coupled to a shank-attached fiber. Light-sensitive ChR2-expressing units were thus depolarized according to the real-time position and orientation of the mouse, generating synthetic spatially selective firing. The instantaneous light intensity administered by the system was calculated by multiplying the maximal illumination by position- and orientation-dependent factors:

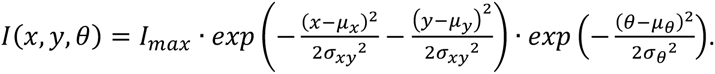

The horizontal center of the two-dimensional position Gaussian, *µ_x_*, was randomized between sessions, drawn from the [25,125] cm range. The horizontal SD of the Gaussian, *α_x_*, was drawn from the [6,20] cm range, controlling the spatial width of illumination (examples in **Fig. S1A**, bottom right). The vertical center, *µ_y_*, was fixed at the vertical center of the track, and the vertical SD, *α_y_*, was the same as *α_x_*. The orientation factor was a one-dimensional Gaussian for which *µ_θ_* was either 0 or π rad depending on the chosen running direction, and *α_θ_* was fixed at 0.87 rad. The maximal illumination level *I_max_* was determined at the beginning of every session during the baseline recordings, chosen to induce robust spiking while avoiding the induction of high-frequency oscillations (Stark et al., 2014). The actual median [IQR] *I_max_* employed was 0.92 [0.51 1.86] µW in CA1 (n=69 sessions) and 6.36 [2.25 15.41] µW in the PPC (n=22 sessions). During every session, illumination was administered during alternate “Light” trials at a single running direction (i.e., once every four trials; **Fig. 2A**). The alternating same-direction trials without illumination are denoted as “Control” trials, and trials in the non-illuminated direction are denoted as “Return” trials. No illumination was given during the first ten trials in either direction.

### Spike detection and sorting

Neural activity was filtered, amplified, multiplexed, and digitized on the headstage (0.1-7,500 Hz, x192; 16 bits, 20 kHz; RHD2132 or RHD2164, Intan Technologies), and then recorded by an RHD2000 evaluation board (Intan Technologies). Offline, spikes were detected and sorted into single units automatically using KlustaKwik3 (Rossant et al., 2016) for shanks with up to 11 sites/shank, or KiloSort2 (Pachitariu et al., 2016) for 16-channel shanks. Automatic spike sorting was followed by manual adjustment of the clusters. Only well-isolated units were used for further analyses (amplitude >40 µV; L-ratio <0.05; ISI index <0.2; (Levi et al., 2022). Units were classified into putative PYR or PV-like INT using a Gaussian mixture model as previously described (Stark et al., 2013).

### Linear track firing rates and place field detection

Generation of trial-averaged firing rate maps and detection of place fields were carried out as previously described (Sloin et al., 2022b). Briefly, the horizontal position on the linear track was binned into 2.5 cm bins, and a firing rate map was created for each unit by dividing the spike count map by the temporal occupancy map. Trials in which the mean running speed was below 10 cm/s were excluded from analyses. Units that fired at least 5 spikes in at least one 2.5 cm spatial bin, pooled over all Light trials, were considered active. Units with significant correlation between the firing rate maps of all Light trial pairs were considered stable. Spontaneous place fields were defined as regions in which the Poisson probability to obtain the observed firing rates or higher was smaller than expected by chance, determined by the baseline on-track firing rate. In CA1, 1,095/4,554 (24%) of the recorded PYRs and 158/716 (22%) of the recorded INTs were active and stable during Light trials. In PPC, 195/360 (54%) of the recorded PYRs were active and stable during Light trials.

### Illumination-dependent firing rate responses

Light limits were defined as the left-most and the right-most bins in which the mean power during Light trials was above three SDs from the trial-averaged mean power during Control trials.

The light-dependent firing rate gain was defined as the mean firing rate within light limits during Light trials, divided by the mean firing rate within light limits during Control trials.

Light-modulated units were defined as units with significant rank correlation (p<0.05, permutation test) between illumination envelope and the difference in mean firing rate during Light and Control trials.

Units were categorized into one of four categories (**Fig. 2D**). Light-modulated units with a firing rate gain larger than one were categorized as “Spatially amplified” if active within light limits during Control trials (n>30 spikes pooled over all trials), and as “Artificial fields” otherwise. Light-modulated units with a firing rate gain smaller than one were categorized as “Rate decreasing” units. All other units were categorized as “Unchanged”.

“Field selectivity” was defined as the mean firing rate within Light limits divided by the mean firing rate outside light limits and was calculated separately for Light and Control trials.

### Prediction of artificial fields

We used cross-validated classifiers (support vector machines, SVMs) to predict the generation of artificial fields during Light trials. We trained n=100 SVMs using ten-fold cross-validation and compared the performance to classifiers trained in the same manner but with shuffled labels. The features used as input were baseline firing rate on the track and spatial information during Control and Return trials, within-limits firing rate during Control trials, field specificity during Control trials, light power, and light intensity (**Fig. S5B-G**). The labelled dataset was comprised of 286 PYRs with artificial fields and 809 without artificial fields, corresponding to chance level of 74%. The accuracy of the classifiers trained in the original dataset was 93%, higher than the accuracy of shuffled labels classifiers (74%; p=7.7×10^-64^, U-test; **Fig. S5H**).

### Theta phase and spatial theta phase precession

To account for the asymmetric shape of LFP theta oscillations, the theta phase at each time point was calculated using the waveform method as previously described (Belluscio et al., 2012; Sloin et al., 2022b). Briefly, in each session, the channel with the best theta signal-to-noise ratio was determined, and the wideband signal from that channel was filtered with a zero-phase two-pole Butterworth band-pass (1-60 Hz). Phases were then defined by linear interpolation between the local maxima (0 rad) and minima (π rad) of the filtered signal.

Spatial precession was quantified by a circular-linear model (Schmidt et al., 2009; Sloin et al., 2022b). To quantify theta phase changes, we fitted a line to the instantaneous theta phase of each spike, *θ(x)*, and the spatial position of the animal at spike time, *x:*

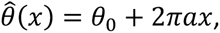

where *a* is the slope and *θ_0_* is the phase offset. For each slope in the tested range ([−0.1, −0.001]), we first calculated the angular residuals of the circular-linear model:

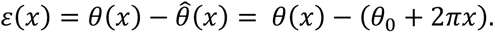

The fitted slope *a* was the slope that yielded the largest *R*, referred to as *R_0_*, corresponding to the slope for which the variance of the residuals was the smallest. The value of the maximal resultant length was used to indicate the fit of the circular-linear model. Statistical significance of the optimized model was estimated by a permutation test, randomly shuffling the {*8(x)*,*x*} pairs for 300 repetitions (Sloin et al., 2022b). The fit of the shuffled phase-position pairs to a circular-linear model was calculated, generating a distribution of fits, *R_p_*. If the probability of observing an *R* value larger than or equal to *R_0_* was smaller than the significance threshold, the field was considered to undergo spatial precession.

Spatial precession effect size η*_s_* was defined as

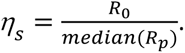

### Constrained randomization test for detecting temporal theta phase precession

The frequency of spikes relative to the LFP was estimated by calculating the spectrum of the cumulative theta phase of the spike train. The frequency at which the spectrum reaches maximal magnitude was defined as the peak frequency of spikes relative to LFP (Mizuseki et al., 2009). A peak frequency of one indicates a fixed 1:1 relation between the frequency of spikes and LFP, i.e., theta phase lock. A peak frequency above one indicates that the instantaneous periodicity of the spikes is faster than the LFP periodicity, i.e., precession.

To detect temporal precession, previous studies (Eliav et al., 2018; Qasim et al., 2021) compared the peak of the cumulative spike phase spectrum between the original and randomized spike trains. Spike phase randomization was performed by drawing phases from the empirical distribution of cumulative LFP phases within each cycle. When phases are generated by a Hilbert transform of the theta-band LFP, the process is equivalent to drawing the phases from a uniform distribution between 0 and 2rc. The spectrum of the phase-randomized trains was then calculated to generate a null distribution of spectra. Although uniform spike-phase randomization can successfully identify units undergoing temporal precession (**Fig. S2A**), the previously employed procedures are prone to two types of errors. First, false positives may occur in phase-locked units, which exhibit a peak around but not necessarily exactly at one (**Fig. S2B**). Some of the false positives can be avoided by requiring the peak frequency to be above one. However, in cases where the observed peak deviates only slightly from one, phase-locked units are still mistakenly classified as undergoing precession. Second, false negatives can occur when a phase-locked unit also exhibits precession, evident as a bimodal cumulative spike phase spectrum (**Fig. S2C**). Consequently, the previously employed randomization approaches are sensitive to false positives and cannot detect precession when the phase-locking peak exhibits a higher magnitude than the precession peak.

We therefore developed a novel constrained randomization test based on the marginal distribution of the spiking phase. In the new approach, random phases were drawn from the empirical distribution of spiking rate as a function of theta phase, as opposed to the empirical distribution of LFP phases or a uniform distribution (**Fig. S2D**). Spike trains randomized under the spiking phase marginal distribution necessarily retain all first-order properties of that distribution. In contrast, the conditional (joint) distribution of spikes and theta phases is not maintained, resulting in the loss of phase changes (e.g., precession; **Fig. S2E**).

The full procedure of the spiking phase constrained randomization test is described in a flowchart (**Fig. S2F**). At the initial stage, the spectrum of the cumulative phase train was estimated, and randomization constrained by LFP phases was employed to identify any significant peaks. Randomization by LFP was carried out at this stage to allow detecting both phase locking and precession. If a significant peak was observed, further assessment was conducted to determine if precession occurred. Phases were redrawn from the empirical spiking phase distribution, and the phase spectrum was recalculated (n=300 repetitions). Subsequently, the mean spiking phase randomized spectrum was subtracted from the original spectrum. The frequency of the peak of the difference between the spectra is denoted as *f_Δ_*. If *f_Δ_* was a local maximum in the original spectrum and was larger than one, the spike train was considered a candidate for temporal precession. Next, the probability of observing temporal precession by chance was estimated by comparing the magnitude of the original spectrum at *f_Δ_*, *m_Δ_*, to the distribution of magnitudes obtained from the randomized spectra, *m_ΔR_*. Since random phases were drawn from a finite pool and since randomization was carried out independently for every spike, multiple spikes could receive the same phase. To account for the effect of spike decimation on spectrum magnitude, the original spike train was subsampled to contain the minimal number of spikes following randomization. If the probability of observing *m_Δ_* (or a higher magnitude) under empirical randomization was smaller than the significance threshold α, the original spike train was classified as undergoing temporal precession.

The temporal precession effect size η_t_ was defined as

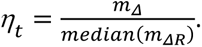

### Half-field analyses

To estimate the consistency of precession slope and effect size across a place field, precessing fields were split into two halves: until the peak mean firing rate, and from the peak firing rate. All (341/341) fields with spontaneous spatial precession and 35/49 (71%) of the artificial fields with synthetic spatial precession exhibited more than 10 spikes in each of the halves, qualifying for the half-field analyses. In every field, precession slope and effect size were estimated separately for each half. Place field symmetry was quantified by the base-10 logarithm of the ratio between the size of the second half-field and the size of the first half-field.

### Quantification of light-induced reduction in precession prevalence

To determine whether the ratio of precession prevalence during Light compared with Control trials differs from chance level, chance level was derived from spontaneous PYR place fields recorded during Control or Return trials. For every spontaneous place field, the available odd and even trials were allocated to different groups. Of the 531 spontaneous place fields with precession in either the spatial or temporal domain (**Fig. S3A**), 525 (99%) had more than 10 spikes in both groups, qualifying for assessing precession prevalence in both groups. In the spatial domain, the first (“odd” trials) group exhibited precession in 279 fields, the second (“even trials”) group exhibited precession in 278 fields, and 199/278.5 (71%) fields exhibited precession in both groups. In the temporal domain, 316 and 310 fields exhibited precession in the odd and even groups, and 222/313 (71%) exhibited precession in both groups. Taking either domain into account, 413 and 402 spontaneous fields precessed during odd and even trials, and 338 (82%) precessed in both groups of trials. The probability that the ratio of precession differed from chance level (**Fig. 3C**) was then estimated by a goodness-of-fit G-test.

### Model predictions

We considered five models for the generation of precession (Maurer and McNaughton, 2007; Jaramillo and Kempter, 2017; Drieu and Zugaro, 2019) and assessed their predictions for an imposed artificial field. Models were implemented using stochastic point processes with a bin size of 0.001 s as follows. In all panels, theta oscillations were defined as 8.5 Hz sine waves. In all spontaneous and artificial fields (except in **Fig. 1B**, left), spatially dependent excitatory input was a Gaussian centered at 1 s with an SD of 0.22 s. Spike probability depended on the sum of external inputs, and a spike was followed by an absolute refractory period of 70 ms. Inheritance of precession precession (Yamaguchi, 2003; Zugaro et al., 2005; Burgess et al., 2007; Hafting et al., 2008; Jaramillo et al., 2014) was generated with unit probability by pulses of excitatory inputs (width, 0.005 s) separated by 0.0066 s, corresponding to precession of 0.056 cycles/cycle (**Fig. 1A**, left). Dual-input (Chance, 2012; Fernández-Ruiz et al., 2017) was generated by the summation of two Gaussians, one centered at 0.9 s and another centered at 1.1 s, both with SD of 0.22 s (**Fig. 1B**). When input from the first Gaussian was larger, spike phase was drawn from a distribution centered at 1.5π; when inputs from the second Gaussian were larger, phase was drawn from a distribution centered at π/2; and when two inputs were equal, the phase was drawn from a distribution centered at π. All distributions had an SD of π/4. Spreading activation (Jensen and Lisman, 1996; Tsodyks et al., 1996) was generated by the addition of excitatory inputs in each theta cycle (**Fig. 1C**). The first excitatory input (width, 0.005 s) arrived at the peak (2π) of the first theta cycle. Subsequent inputs arrived 0.0066 s (corresponding to 0.056 cycles/cycle) on each consecutive theta cycle, representing shorter cumulative transmission delays from upstream neurons with closer place fields. Inhibition-excitation summation (Kamondi et al., 1998; Magee, 2001; Harris et al., 2002; Mehta et al., 2002) was generated by the summation of the excitatory Gaussian input with inhibitory currents opposite in phase to the ongoing theta (**Fig. 1D**). Finally, dual-oscillator (O’Keefe and Recce, 1993; Bose et al., 2000; Bose and Recce, 2001; Lengyel et al., 2003) was generated by multiplying a 9.1 Hz sinusoid by the input envelope (**Fig. 1E**). In panels **ABC**, spike probability in artificial fields depended only on the excitatory inputs, leading to no theta phase preference. In the other panels, spike probability depended on the difference between excitation and inhibition (**Fig. 1D**, right) and on an excitation-triggered fast oscillator (**Fig. 1E**, right), predicting precession. Similar predictions were obtained using integrate and fire spiking models, and when pooling data over multiple virtual runs along the track with randomized theta onset phases.

### Statistical analyses

For all statistical tests, a significance threshold of α=0.05 was used. For testing the prevalence of either temporal or spatial precession, the threshold was corrected to 1-(1-α)^2^=0.0975. All descriptive statistics (n, mean, median, SD, SEM, IQR, range) can be found in the results, figures, figure legends, and tables. Non-parametric statistical tests were used for all analyses. A one-sample Wilcoxon signed-rank test (two-tailed, unless noted otherwise) was used to determine whether a group median differed from a fixed value (e.g., zero, one). Differences between the medians of two unpaired groups were tested with a two-tailed Mann-Whitney U-test. Differences between the medians of two paired groups were tested with Wilcoxon’s two-sample signed-rank test (two-tailed). Differences between medians of three or more groups were tested with the Kruskal-Wallis test and corrected for multiple comparisons using Tukey’s procedure. An exact one-tailed binomial test was used to estimate whether a given fraction was larger than expected by chance. Differences between the proportions of observations of two categorical variables were tested with a likelihood ratio (G-) test of independence. Differences from expected proportions were tested with a goodness-of-fit G-test. Bonferroni’s correction was employed in case of G-test multiple comparisons. The significance of Spearman’s (rank) correlation coefficients was tested by a permutation test.

## Tables

**Table S1.** Experimental animals.

| Animal ID | Sex | Strain<sup>1</sup> | Age<sup>2</sup><br>[week] | Weight<sup>2</sup><br>[g] | Probe | Optics<sup>3</sup> | CA1 |  | PPC |  |
| --- | --- | --- | --- | --- | --- | --- | --- | --- | --- | --- |
|  |  |  |  |  |  |  | Sessions | PYRs | Sessions | PYRs |
| mA234 | M | FVB x<br>CaMKII::ChR2 | 16 | 30 | Buzsaki32 | 3 x 470<br>nm | 20 | 1,229 | 3 | 70 |
| mC41 | M | FVB x<br>CaMKII::ChR2 | 10 | 33.7 | Stark64 | 5 x 470<br>nm<br>1 x 356<br>nm | 23 | 1,380 |  |  |
| mDS1 | M | CaMKII::ChR2 | 14 | 25.7 | DS64<sup>4</sup> | 2 x 470<br>nm | 5 | 222 |  |  |
| mDS2 | F | CaMKII::ChR2 | 30 | 24.2 | DS64 | 2 x 470<br>nm | 21 | 1,732 |  |  |
| mC89 | M | CaMKII::ChR2 | 19 | 29.7 | DS64 | 2 x 470<br>nm |  |  | 19 | 290 |
| **Total** |  |  |  |  |  |  | **69** | **4,554** | **22** | **360** |

<sup>1</sup> CaMKII::ChR2 mice are the heterozygous offspring of homozygous CaMKII-Cre males (#005359, Jackson Labs) and Ai32 females, homozygous for the Rosa-CAG-LSLChR2(H134R)-EYFP-WPRE conditional allele (JAX #012569). FVB x CaMKII::ChR2 mice are offspring of heterozygous CaMKII::ChR2 males and FVB/NJ females (JAX #001800).
<sup>2</sup> At the time of implantation.
<sup>3</sup> Number of optical fibers coupled to recording shanks x the emitted wavelength.
<sup>4</sup> Dual-sided 64-channel probe.

**Table S2.** Light response and theta phase precession of CA1 PYRs in every mouse.

| Animal ID | Sessions | Trials<sup>1</sup> | Active PYRs<sup>2</sup> | Fields | Spontaneous precession<sup>3</sup> | Light responsive<sup>4</sup> | Spatially amplified<sup>5</sup> | Artificial fields<sup>6</sup> | Synthetic precession<sup>7</sup> |
| --- | --- | --- | --- | --- | --- | --- | --- | --- | --- |
| mA234 | 23 | 193<br>[168 239] | 241 | 139 | 116 | 119 | 79 | 40 | 21 (52%)*** |
| mC41 | 20 | 171<br>[136 206] | 282 | 150 | 122 | 180 | 103 | 77 | 29 (38%)*** |
| mDS1 | 5 | 136<br>[120 140] | 41 | 19 | 11 | 33 | 8 | 25 | 6 (24%)** |
| mDS2 | 21 | 161<br>[131 182] | 531 | 364 | 282 | 344 | 200 | 144 | 49 (34%)*** |
| **Total** | **69** | **168**<br>**[135 200]** | **1,095** | **672** | **531** | **676** | **390** | **286** | **105 (37%)\*\*\*** |

<sup>1</sup> Median [IQR] number of one-direction trials.
<sup>2</sup> PYRs that fired in an active and stable manner during Light trials.
<sup>3</sup> Fields with spatial or temporal precession.
<sup>4</sup> PYRs that exhibited a firing rate increase during light stimulation ( $p < 0.05$ , permutation test).
<sup>5</sup> PYRs that were active within light limits during Control trials and exhibited firing rate increase during Light trials.
<sup>6</sup> PYRs that were not active within light limits during Control trials but were active during Light trials.
<sup>7</sup> PYRs with artificial fields that exhibited precession during Light trials. \*\*/\*\*\*: $p < 0.01/p < 0.001$ , binomial test comparing to chance level (0.0975).

**Table S3.** Correlation of spontaneous and synthetic precession with single unit features.

|  |  | Spontaneous |  |  |  | Synthetic |  |  |  |
| --- | --- | --- | --- | --- | --- | --- | --- | --- | --- |
| Feature class | Feature | cc±SEM<sup>1</sup> | cc p-value<sup>2</sup> | pcc±SEM<sup>3</sup> | pcc p-value<sup>4</sup> | cc±SEM | cc p-value | pcc±SEM | pcc p-value |
| Behavioral features | Number of trials<sup>5</sup> | 0.053±0.0036 | 0.18 | 0.041±0.0038 | 0.31 | -0.037±0.0062 | 0.51 | 0.01±0.0061 | 0.86 |
|  | In-field speed [cm/s]<sup>6</sup> | 0.014±0.0034 | 0.73 | 0.1±0.0044 | 0.006 | 0.067±0.0055 | 0.26 | -0.004±0.0067 | 0.95 |
|  | Time spent in field [s]<sup>6</sup> | 0.015±0.0038 | 0.68 | 0.13±0.0041 | **0.002** | 0.13±0.0058 | 0.036 | 0.085±0.0063 | 0.16 |
| Spiking features | Spike width [ms]<sup>5</sup> | 0.095±0.0044 | 0.012 | 0.051±0.0042 | 0.2 | 0.044±0.0055 | 0.45 | 0.057±0.0058 | 0.36 |
|  | Burst index<sup>5</sup> | 0.04±0.0041 | 0.31 | 0.053±0.0039 | 0.19 | 0.12±0.0057 | 0.039 | 0.13±0.0058 | 0.039 |
| Network features | Post-synaptic units<sup>5</sup> | 0.015±0.0037 | 0.69 | 0.016±0.0045 | 0.69 | -0.033±0.0056 | 0.59 | 0.00027±0.0065 | 1 |
|  | Pre-synaptic INT<sup>5</sup> | 0.064±0.0034 | 0.11 | 0.058±0.0038 | 0.14 | -0.0043±0.0055 | 0.94 | -0.0099±0.0058 | 0.87 |
| On track spiking features | Baseline firing rate [spk/s]<sup>5</sup> | -0.013±0.0037 | 0.72 | -0.11±0.0035 | 0.006 | 0.17±0.0058 | 0.0067 | 0.2±0.0058 | **0.00067** |
|  | SI<sup>6</sup> | 0.19±0.0039 | 0.00067 | 0.14±0.0045 | **0.002** | 0.12±0.0058 | 0.056 | -0.007±0.0067 | 0.91 |
|  | Field size [cm]<sup>6</sup> | -0.037±0.0038 | 0.33 | -0.17±0.0042 | **0.00067** | 0.12±0.005 | 0.042 | 0.038±0.0056 | 0.53 |
|  | Peak in-field firing rate [spk/s]<sup>6</sup> | 0.16±0.0043 | 0.00067 | 0.032±0.0047 | 0.39 | 0.14±0.0057 | 0.029 | 0.043±0.0068 | 0.47 |
| In-field spike-LFP features | Spike lock to theta phase (R)<sup>6</sup> | -0.071±0.0045 | 0.07 | -0.03±0.0041 | 0.45 | -0.00087±0.0072 | 0.99 | 0.065±0.0073 | 0.26 |
|  | Precession slope<sup>6</sup> | -0.2±0.0043 | 0.00067 | -0.22±0.0041 | **0.00067** | -0.091±0.0052 | 0.14 | -0.11±0.0055 | 0.078 |

<sup>1</sup> Rank correlation coefficient.<sup>2</sup> Rank correlation permutation test.<sup>3</sup> Partial rank correlation coefficient.
<sup>4</sup> Permutation test of the partial correlation. Features that exhibit significant correlation after Bonferroni correction for multiple comparisons (i.e., p-value smaller than 0.05/13) are highlighted in bold. P-values for cc and pcc are uncorrected. Together, the 13 features explain 14% of the variance among fields with and without spontaneous precession ( $R^2=0.14$ ; $p=0.00067$ , permutation test) and 11% of the variance among units with and without synthetic precession ( $R^2=0.11$ ; $p=0.0087$ , permutation test).
<sup>5</sup> Unit dependent features, common to both spontaneous and induced fields.<sup>6</sup> Spontaneous/synthetic field specific features.

**Table S4.** Light response and theta phase precession of local CA1 INTs in every mouse.

| Animal | Sessions | Trials | Active INTs | Fields | Spontaneous precession | Light responsive | Spatially amplified | Artificial fields | Synthetic precession |
| --- | --- | --- | --- | --- | --- | --- | --- | --- | --- |
| mA234 | 23 | 193<br>[168 239] | 37 | 12 | 2 | 5 | 4 | 1 | 1<sup>ns</sup> |
| mC41 | 20 | 171<br>[136 206] | 29 | 16 | 9 | 19 | 18 | 1 | 0 |
| mDS1 | 5 | 136<br>[120 140] | 6 | 1 | 0 | 4 | 4 | 0 | 0 |
| mDS2 | 21 | 161<br>[131 182] | 86 | 31 | 7 | 57 | 56 | 1 | 0 |
| **Total** | **69** | **168**<br>**[135 200]** | **158** | **60** | **18** | **85** | **82** | **3** | **1<sup>ns</sup>** |

All conventions are the same as in **Table S2**.

**Table S5.** Light response and theta phase precession of PPC PYRs in every mouse.

| Animal | Sessions | Trials | Active PYRs | Fields | Spontaneous precession | Light responsive | Spatially amplified | Artificial fields | Synthetic precession |
| --- | --- | --- | --- | --- | --- | --- | --- | --- | --- |
| mA234 | 3 | 99<br>[68 163] | 25 | 1 | 0 | 23 | 8 | 15 | 1 (7%)<sup>ns</sup> |
| m89C | 19 | 146<br>[118 167] | 170 | 36 | 6 | 136 | 99 | 37 | 3 (8%)<sup>ns</sup> |
| **Total** | **22** | **146**<br>**[110 168]** | **195** | **37** | **6** | **159** | **107** | **52** | **4 (8%)<sup>ns</sup>** |

All conventions are the same as in **Table S2**.

## Supplementary figures

**Figure S1.**
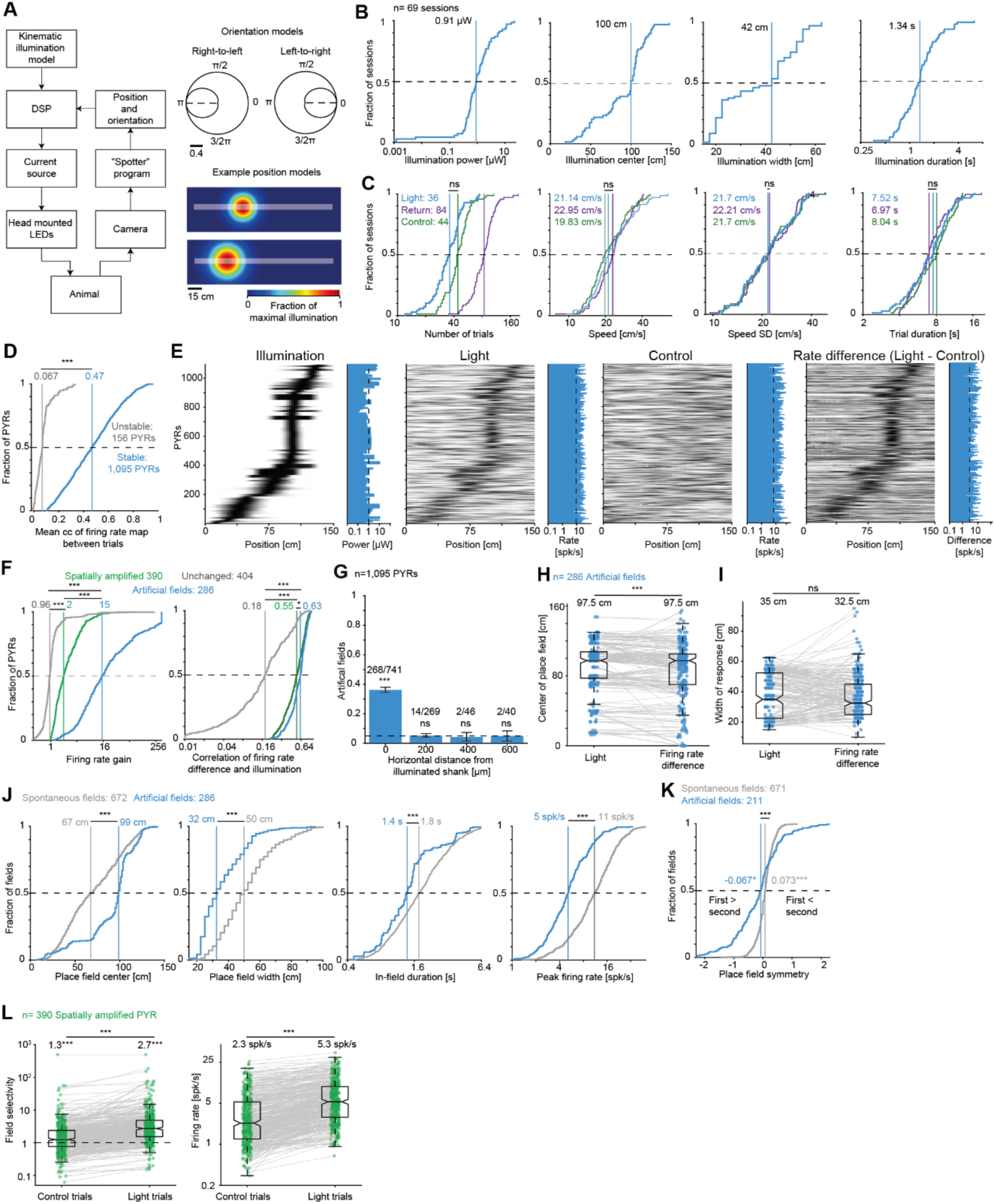
CA1 activation produces artificial place fields. (**A**) Procedure for closed-loop orientation- and position-dependent illumination. Left, System components. Right, Example kinematic models. (**B**) Illumination characteristics: Maximal power (*I_max_*), center (*µ_x_*), width (*σ_x_*), and duration. Here and in **CDFJL**, vertical lines show group medians. (**C**) Illumination does not affect running speed. Parameters during Light, Control, and Return trials: number of trials, trial-averaged speed, speed SD, trial duration. Here and in **F**, ns/***: p>0.05/p<0.001, Kruskal-Wallis test, corrected for multiple comparisons. (**D**) PYR firing rate maps are stable during Light trials. The stability of each recorded PYR was estimated by the mean rank correlation coefficient (cc) between the firing rate maps of all pairs of Light trials. Here and in **JL**, lined ***: p<0.001, U-test. (**E**) Position-dependent illumination generates an increase in firing rates in matching regions. Each row represents a single CA1 PYR, sorted according to the position at which illumination peaked. Histograms represent the peak of each unit. (**F**) Firing rate gain (left) and correlation between firing rate and illumination (right) differ between the PYRs of the four activation groups. (**G**) Artificial fields occur locally, on the illuminated shank. Horizontal dashed line shows chance level; error bars, SEM. ns/***: p>0.05/p<0.001, G-test. (**H**) Artificial place fields peak slightly before illumination peaks. Here and in **IL**, lined ns/***: p>0.05/p<0.001, Wilcoxon’s test. (**I**) Width of illumination matched with the width of each artificial place field. (**J**) Properties of artificial fields during Light trials, compared with properties of spontaneous place fields recorded during Control and Return trials. (**K**) Place field symmetry, quantified by the base-10 logarithm of the ratio between the size of the second half-field and the size of the first half-field. Artificial fields are skewed to the left (larger first half), whereas spontaneous fields are skewed to the right (larger second half). Only fields with at least 10 spikes in both halves are included. */***: p<0.05/p<0.001, Wilcoxon’s test compared with a zero null. (**L**) Light increases the firing rates of PYRs with preexisting place fields. Left, Field selectivity of spatially amplified PYRs is above one during Light and Control trials. ***: p<0.001, Wilcoxon’s test compared to a unity null. Right, Mean firing rate within light limits is higher during Light compared with Control trials.

**Figure S2.**
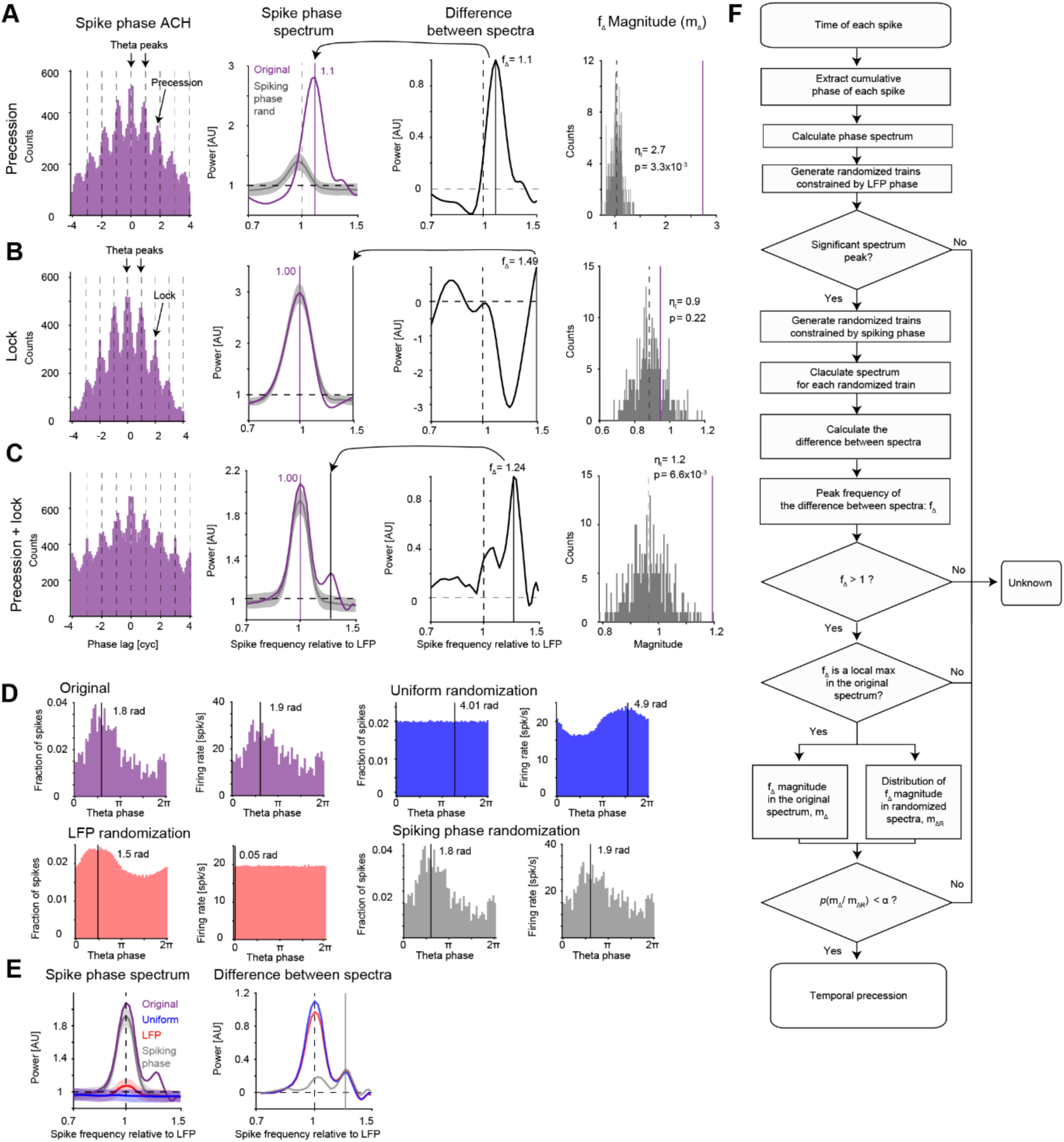
Constrained randomization test for detecting temporal theta phase precession. (**A**) Demonstration of the spiking phase constrained randomization test for a phase precessing unit. First column, ACH of the cumulative theta phase of each spike. Dashed lines show theta cycle peaks. ACH peaks occur before theta peaks, suggesting temporal precession. Second column, The spectrum of the PYR spike train transformed to cumulative phase (purple), and of spike trains randomly drawn from the marginal distribution of spiking phases (gray; mean±SEM, 300 repetitions). Third column, The difference between the original spike phase spectrum and the spectra of spike phase trains randomized from the marginal spike phase distribution. The frequency with the maximal positive difference in magnitude is denoted *f_Δ_*. Fourth column, Comparison of the spectral power magnitude at *f_Δ_* in the original spike phase spectrum (purple) versus the magnitude at *f_Δ_* for the randomized spectra (gray). (**B**) Demonstration of the constrained spiking phase randomization test for a phase locked PYR. (**C**) Demonstration of the test for a PYR expressing both phase lock and precession. (**D**) Original phase distribution of the example unit in **C** (top left), and phases randomized under three different constraints. In every case, the phase of each spike was drawn randomly within the specific theta cycle. Spike phases can be drawn from one of three distributions. (1) Uniform between 0 and 2π (“Uniform randomization”). (2) The marginal distribution of the theta wave phases (“LFP randomization”). (3) The marginal distribution of spike phases (“Spiking phase randomization”). Left, The fraction of spikes at every theta phase. Right, Firing rate at every phase. By definition, only the spiking phase randomization method replicates the marginal phase distribution of the original spike train. (**E**) The spectrum of the original cumulative phase spike train and cumulative randomized phases. Only randomization under the spiking phase distribution constraint allows disambiguating and identifying precession in the presence of phase lock. (**F**) Flowchart describing the steps of the spiking phase constrained randomization test for detecting precession. Quantifying the cumulative phase spike spectrum combined with the constrained randomization allows the statistical identification of temporal precession.

**Figure S3.**
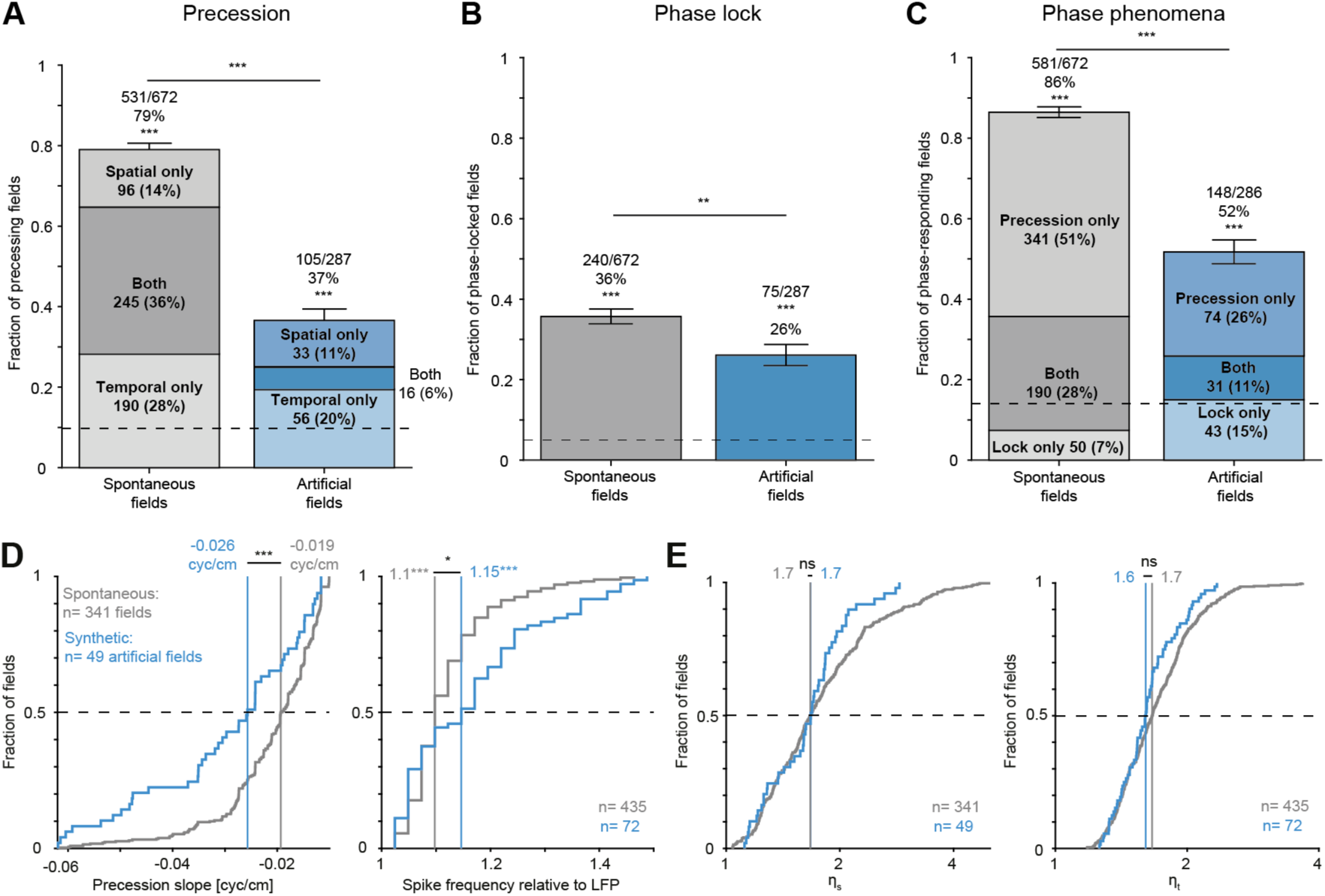
PYR activation induces synthetic precession and phase locking. (**A**) Artificial field spikes exhibit synthetic precession. Here and in **BC**, ***: p<0.001, binomial test. Lined **/***: p<0.001/p<0.001, G*-*test. (**B**) Artificial field spikes exhibit synthetic phase lock. (**C**) More than half of the artificial fields exhibit either synthetic theta phase lock or precession. Horizontal dashed line, chance level for detecting any theta phase phenomenon, 1-(1-α)^3^=0.14. (**D**) Synthetic precession is faster than spontaneous precession. Left, Synthetic precession exhibits a steeper spatial slope (median [IQR]: −0.026 [−0.04 −0.02] cyc/cm) compared with spontaneous precession (−0.019 [−0.03 −0.01] cyc/cm; ***: p=1.7×10^-4^, U-test). Right, Synthetic temporal precession shows higher spike frequency relative to LFP (1.15 [1.05 1.24]) compared with spontaneous precession (1.1 [1.07 1.15]; *: p=0.046). (**E**) Left, The effect size of spatial precession is 1.67 [1.36 2.18] for spontaneous precession, compared to 1.67 [1.35 1.88] for synthetic precession (ns: p=0.33, U-test). Right, The effect size of temporal precession is 1.66 [1.45 1.91] for spontaneous precession, compared with 1.61 [1.44 1.77] for spontaneous precession (ns: p=0.11).

**Figure S4.**
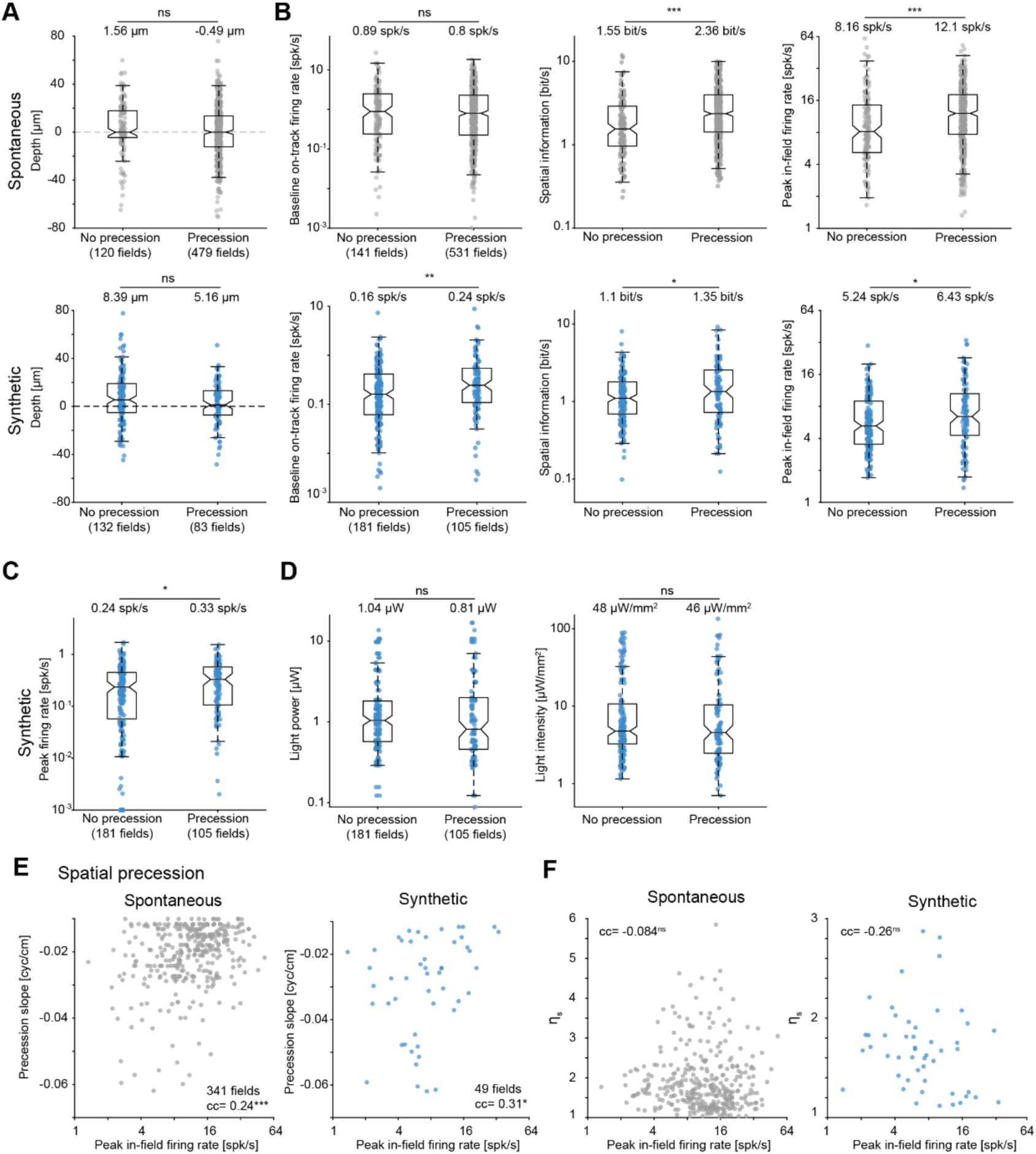
Spontaneous and synthetic precession correlate with firing rates. (**A**) Spontaneous (top) and synthetic (bottom) precession prevalence is not associated with soma depth within CA1 str. pyramidale. Only PYRs for which the depth within the layer could be determined are included. Here and in **BCD**: For spontaneous precession, “No precession” PYRs exhibit spontaneous place fields but no precessing fields, whereas “Precession” PYRs exhibit at least one phase-precessing field. For synthetic precession, “No precession” PYRs have non-precessing artificial place fields, whereas “Precession” PYRs have artificial fields with synthetic precession in either the spatial or temporal domain. Ns/*/**/***: p>0.05/p<0.05/p<0.01/p<0.001, U-test. (**B**) Spontaneous and synthetic precession prevalence is correlated with on-track firing rates. Left, Baseline firing rates on track. Middle, Spatial information. Right, Peak in-field firing rates. (**C**) The firing rate during Control trials is higher for units with synthetic precession than for units with non-precessing artificial fields. (**D**) Light intensity at the estimated soma location is not different between units with synthetic precession and non-precessing artificial fields. (**E**) Spatial precession slope is correlated with peak in-field firing rates in spontaneous and synthetic precession. Here and in **F**, ns/*/***: p>0.05/p<0.05/p<0.001, permutation test. (**F**) Spatial precession effect size is not correlated with peak firing rates.

**Figure. S5.**
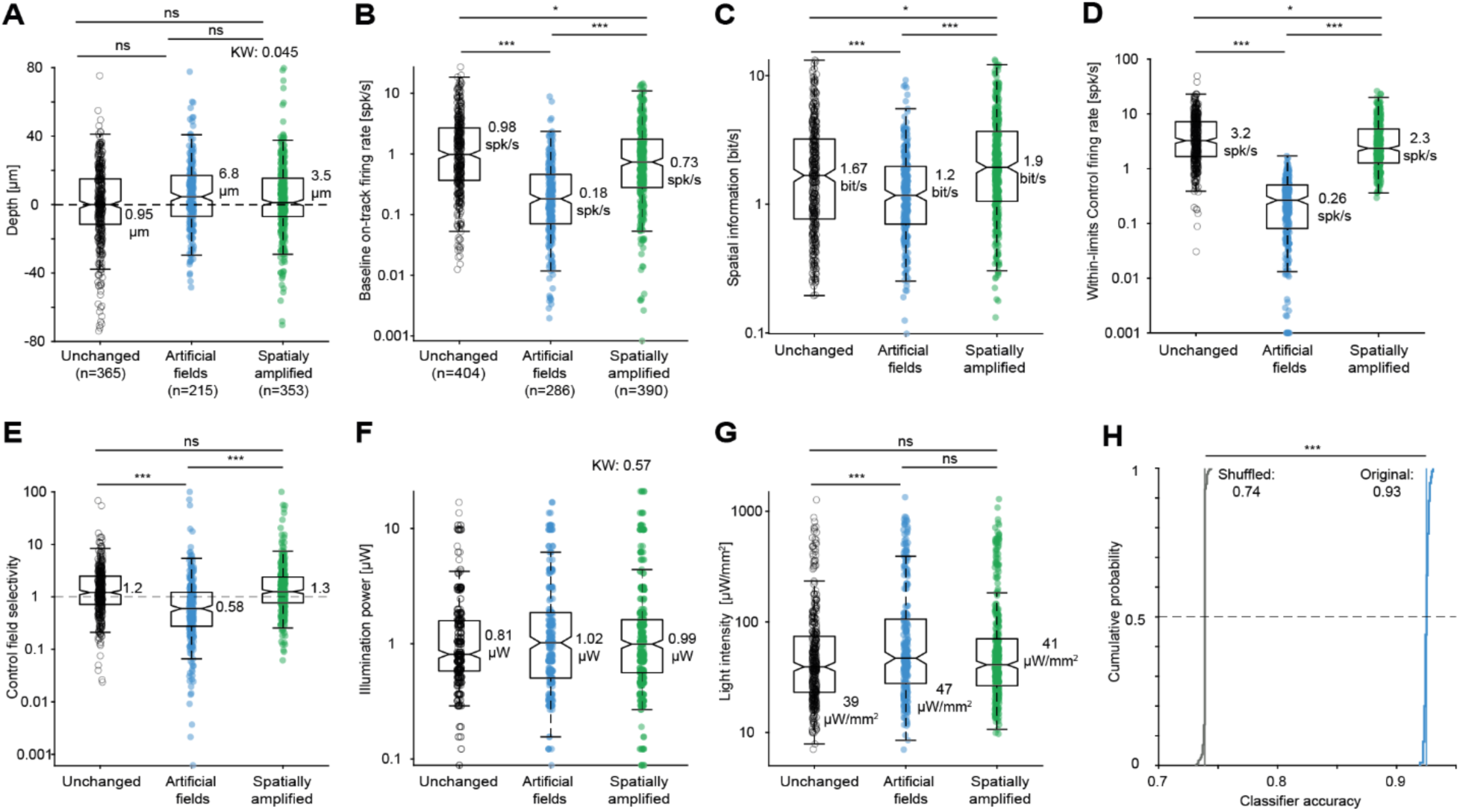
Spiking during Control trials and illumination specifics predict which PYRs exhibit artificial fields. (**A**) The specific light-activation pattern – artificial fields, spatially amplified fields, or unchanged – is associated with soma depth within CA1 str. pyramidale. Only PYRs for which the depth within the layer could be determined are included. Here and in **F**, the p-value in text is for a Kruskal-Wallis test. Here and in **B-G**, ns/*/***: p>0.05/p<0.01/p<0.001, Kruskal-Wallis test corrected for multiple comparisons. (**B**) Light activation pattern is correlated with on-track firing rates during Control trials. (**C**) Light activation pattern is correlated with spatial information. (**D**) During Control trials, PYRs with artificial place fields exhibit lower firing rates within the light limits, compared with other light activation patterns. (**E**) During Control trials, PYRs with artificial fields exhibit lower field specificity for firing within the illuminated region, compared with PYRs that exhibit other light activation patterns. (**F**) Illumination power does not differ between PYRs with and without artificial fields. (**G**) Light intensity at the estimated soma location is higher in PYRs exhibiting artificial fields, compared with no change PYRs. (**H**) Ten-fold cross-validated classifiers (SVM) trained on the features depicted in **B-G** predict which PYRs show artificial place fields (n=100 independent runs). In contrast, classifiers trained with shuffled labels exhibit chance performance (74%). ***: p<0.001, U-test.

**Figure S6.**
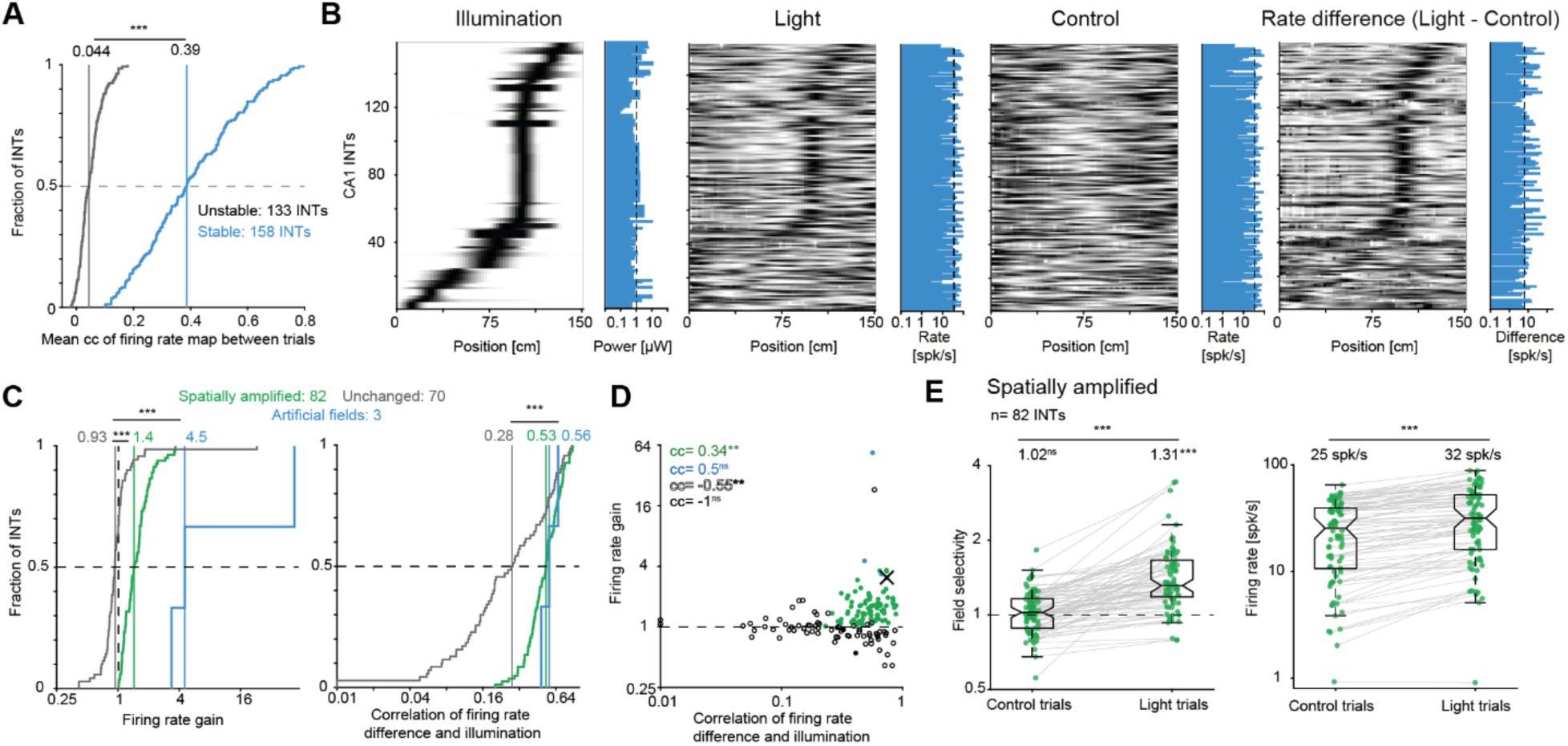
CA1 PYR activation increases the firing rates of spontaneously active INT. (**A**) INT firing rate maps are stable during Light trials. ***: p<0.001, U-test. All conventions are the same as in the corresponding panels of Fig. 1 and **Fig. S1**. (**B**) Closed-loop PYR activation in CA1 generates an indirect increase of INT firing rates. Each row corresponds to a single INT determined to be active and stable during Light trials recorded on the illuminated shank. (**C**) Firing rate increases of CA1 INTs during CA1 PYR activation. Left, Firing rate gain. Right, Correlation between firing rate and illumination. ***: p<0.001, Kruskal-Wallis test. (**D**) Firing rate gain vs. firing rate correlation with light pattern. ns/**: p>0.05/p<*0*.01, permutation test. X, example INT (Fig. 5A). (**E**) Illumination amplifies the firing rate of INTs that exhibit spontaneous spatial selectivity. Left, The field selectivity of spatially amplified INTs is above one during Light but not during control trials. ns/***: p>0.05/p<0.001, Wilcoxon’s test. Right, The mean firing rate within light limits is higher during Light compared with Control trials.

**Figure S7.**
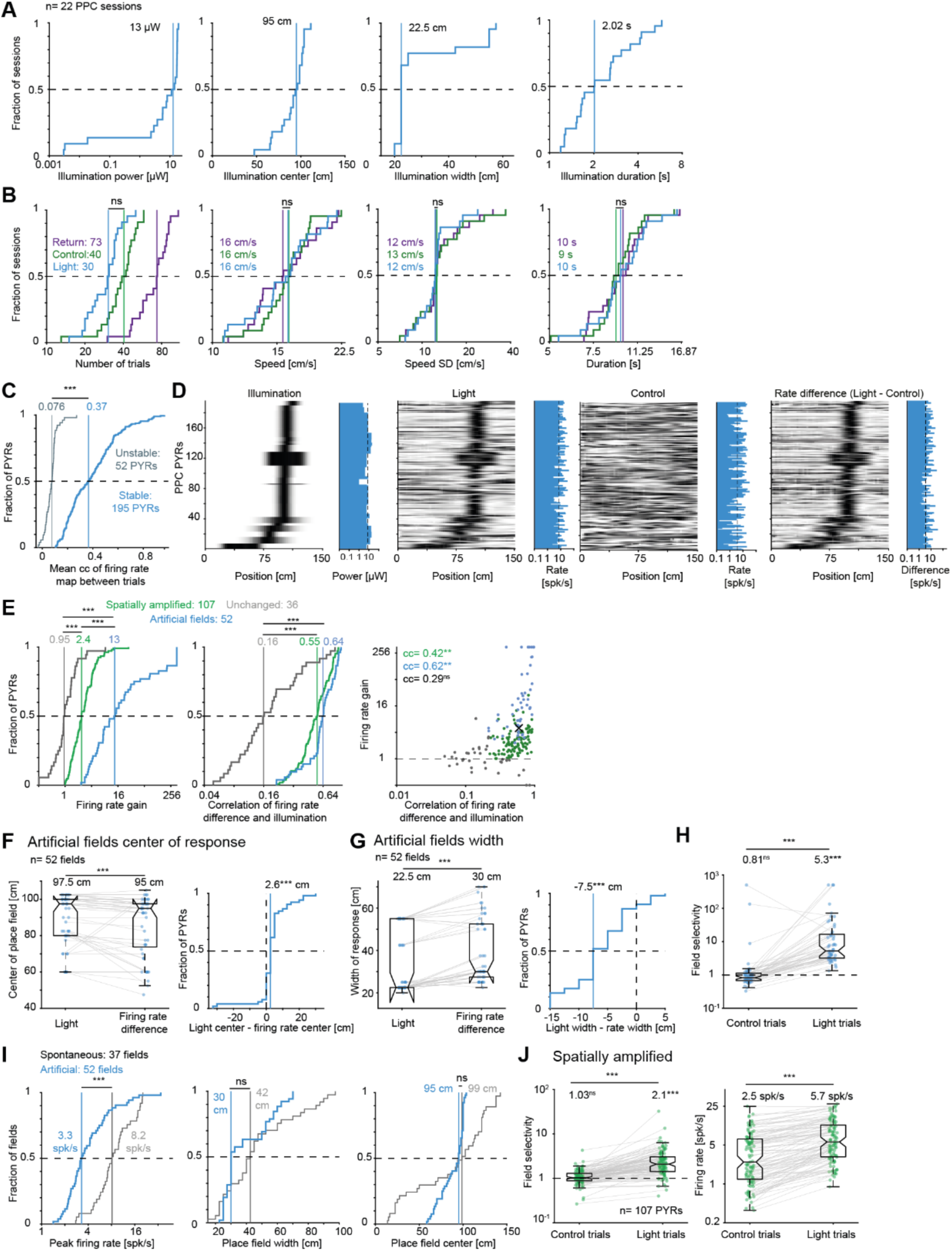
Activation of PPC PYR generates artificial place fields. (**A**) Illumination characteristics: Maximal power, center, width, and duration. All conventions are the same as in the corresponding panels of Fig. 2 and **Fig. S1**. (**B**) Illumination does not affect running speed. Parameters during Light, Return, and Control trials: Number of trials, trial-averaged speed, speed SD, and trial duration. Here and in **E**, ns/***: p>0.05/p<0.001, Kruskal-Wallis test, corrected for multiple comparisons. (**C**) Firing rate maps of PPC PYRs are stable during Light trials. Here and in **HI**, lined ns/***: p>0.05/p<0.001, U-test. (**D**) PPC position-dependent illumination generates an increase in firing rates in matching regions. (**E**) Illumination generates artificial place fields in PPC PYRs. Firing rate gain (left) and correlation between firing rate and illumination (middle) differ between PYRs of the activation groups. Right, Firing rate gain vs. firing rate correlation with light pattern. ns/**: p>0.05/p<0.01, permutation test. X, example PPC PYR (Fig. 5G). (**F**) Artificial place fields peak slightly before illumination peaks. Left, Peak of illumination matched with the peak of each artificial place field. Here and in **HJ**, ns/***: p>0.05/p<0.001, Wilcoxon’s test. Right, The spatial distance between illumination peak and firing rate increase peak. (**G**) Artificial place fields and illumination have smaller spatial widths. Left, Width of illumination matched with the width of each artificial place field. Right, Difference between the width of the artificial field and the received illumination. (**H**) Field selectivity is above one during Light but not during Control trials. (**I**) Properties of artificial fields during Light trials, compared with the properties of spontaneous place fields. (**J**) Light increases the firing rates of PYRs spontaneously active within light limits. Left, The field selectivity of spatially amplified PPC PYRs is above one during Light and Control trials. Right, Mean firing rate within light limits is higher during Light compared with Control trials.

**Figure S8.**
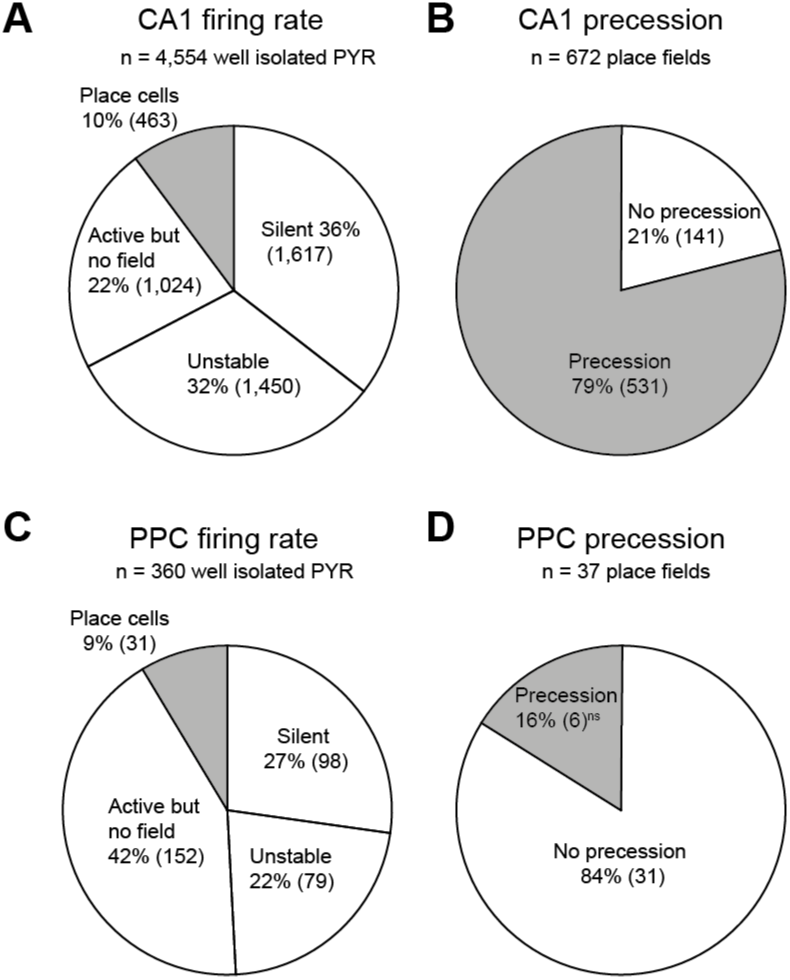
PPC PYRs exhibit spontaneous spatially modulated firing but no spontaneous precession. (**A**) Spontaneous rate coding in CA1 PYRs. “Silent” PYRs fire less than 30 spikes across all trials on the linear track. “Unstable” PYRs do not exhibit correlation between the firing rate maps of all pairs of Control trials. “Active but no field” PYRs fire on the track but do not exhibit a distinct preferred location. “Place cells” exhibit one or more spontaneous place fields. (**B**) The majority of spontaneous CA1 place fields exhibit theta phase precession. (**C**) Spontaneous rate coding in the PPC. PPC PYRs exhibit spatially preferred regions which are defined as place fields, using criteria identical to those used with CA1 PYRs. (**D**) PPC spontaneous place fields do not feature theta phase precession. ns: p=0.15, binomial test comparing to chance level (0.0975).

## References

Belluscio MA, Mizuseki K, Schmidt R, Kempter R, Buzsáki G (2012) Cross-frequency phase–phase coupling between theta and gamma oscillations in the hippocampus. J Neurosci 32:423–435.

Bose A, Booth V, Recce M (2000) A temporal mechanism for generating the phase precession of hippocampal place cells. J Comput Neurosci 9:5–30.

Bose A, Recce M (2001) Phase precession and phase-locking of hippocampal pyramidal cells. Hippocampus 11:204–215.

Burgess N, Barry C, O’Keefe J (2007) An oscillatory interference model of grid cell firing. Hippocampus 17:801–812.

Bush D, Ólafsdóttir HF, Barry C, Burgess N (2022) Ripple band phase precession of place cell firing during replay. Curr Biol 32:64–73.

Chadwick A, van Rossum MC, Nolan MF (2016) Flexible theta sequence compression mediated via phase precessing interneurons. eLife 5:e20349.

Chance FS (2012) Hippocampal phase precession from dual input components. J Neurosci Off J Soc Neurosci 32:16693–16703a.

Dragoi G, Buzsáki G (2006) Temporal encoding of place sequences by hippocampal cell assemblies. Neuron 50:145–157.

Drieu C, Todorova R, Zugaro M (2018) Nested sequences of hippocampal assemblies during behavior support subsequent sleep replay. Science 362:675–679.

Drieu C, Zugaro M (2019) Hippocampal sequences during exploration: Mechanisms and functions. Front Cell Neurosci 13:232.

Ego-Stengel V, Wilson MA (2007) Spatial selectivity and theta phase precession in CA1 interneurons. Hippocampus 17:161–174.

Eliav T, Geva-Sagiv M, Yartsev MM, Finkelstein A, Rubin A, Las L, Ulanovsky N (2018) Nonoscillatory phase coding and synchronization in the bat hippocampal formation. Cell 175:1119–1130.

Fernández-Ruiz A, Oliva A, Nagy GA, Maurer AP, Berényi A, Buzsáki G (2017) Entorhinal-CA3 dual-input control of spike timing in the hippocampus by theta-gamma coupling. Neuron 93:1213–1226.e5.

Foster DJ, Wilson MA (2006) Reverse replay of behavioural sequences in hippocampal place cells during the awake state. Nature 440:680–683.

Gaspar N, Eichler R, Stark E (2019) A novel low-noise movement tracking system with real-time analog output for closed-loop experiments. J Neurosci Methods 318:69–77.

George TM, de Cothi W, Stachenfeld KL, Barry C (2023) Rapid learning of predictive maps with STDP and theta phase precession. eLife 12:e80663.

Grienberger C, Milstein AD, Bittner KC, Romani S, Magee JC (2017) Inhibitory suppression of heterogeneously tuned excitation enhances spatial coding in CA1 place cells. Nat Neurosci 20:417–426.

Hafting T, Fyhn M, Bonnevie T, Moser MB, Moser EI (2008) Hippocampus-independent phase precession in entorhinal grid cells. Nature 453:1248–1252.

Harris KD, Henze DA, Hirase H, Leinekugel X, Dragoi G, Czurkó A, Buzsáki G (2002) Spike train dynamics predicts theta-related phase precession in hippocampal pyramidal cells. Nature 417:738–741.

Harvey CD, Collman F, Dombeck DA, Tank DW (2009) Intracellular dynamics of hippocampal place cells during virtual navigation. Nature 461:941–946.

Hirase H, Czurkó A, Csicsvari J, Buzsáki G (1999) Firing rate and theta-phase coding by hippocampal pyramidal neurons during ‘space clamping.’ Eur J Neurosci 11:4373–4380.

Hutcheon B, Yarom Y (2000) Resonance, oscillation and the intrinsic frequency preferences of neurons. Trends Neurosci 23:216–222.

Huxter J, Burgess N, O’Keefe J (2003) Independent rate and temporal coding in hippocampal pyramidal cells. Nature 425:828–832.

Izhikevich EM (2006) Dynamical Systems in Neuroscience: The Geometry of Excitability and Bursting. MIT press.

Jaramillo J, Kempter R (2017) Phase precession: A neural code underlying episodic memory? Curr Opin Neurobiol 43:130–138.

Jaramillo J, Schmidt R, Kempter R (2014) Modeling inheritance of phase precession in the hippocampal formation. J Neurosci:7715–7731.

Jensen O, Lisman JE (1996) Hippocampal CA3 region predicts memory sequences: Accounting for the phase precession of place cells. Learn Mem 3:279–287.

Jensen O, Lisman JE (2000) Position reconstruction from an ensemble of hippocampal place cells: Contribution of theta phase coding. J Neurophysiol 83:2602–2609.

Jones MW, Wilson MA (2005) Phase precession of medial prefrontal cortical activity relative to the hippocampal theta rhythm. Hippocampus 15:867–873.

Kamondi A, Acsády L, Wang XJ, Buzsáki G (1998) Theta oscillations in somata and dendrites of hippocampal pyramidal cells in vivo: Activity-dependent phase-precession of action potentials. Hippocampus 8:244–261.

Krumin M, Lee JJ, Harris KD, Carandini M (2018) Decision and navigation in mouse parietal cortex. eLife 7:e42583.

Lee D, Lin B-J, Lee AK (2012) Hippocampal Place Fields Emerge upon Single-Cell Manipulation of Excitability During Behavior. Science 337:849–853.

Lengyel M, Szatmáry Z, Érdi P (2003) Dynamically detuned oscillations account for the coupled rate and temporal code of place cell firing. Hippocampus 6:700–714.

Levi A, Spivak L, Sloin HE, Someck S, Stark E (2022) Error correction and improved precision of spike timing in converging cortical networks. Cell Rep 40:111383.

Magee JC (2001) Dendritic mechanisms of phase precession in hippocampal CA1 pyramidal neurons. J Neurophysiol 86:528–532.

Maurer AP, Cowen SL, Burke SN, Barnes CA, McNaughton BL (2006) Phase precession in hippocampal interneurons showing strong functional coupling to individual pyramidal cells. J Neurosci 26:13485–13492.

Maurer AP, Lester AW, Burke SN, Ferng JJ, Barnes CA (2014) Back to the future: preserved hippocampal network activity during reverse ambulation. J Neurosci 34:15022–15031.

Maurer AP, McNaughton BL (2007) Network and intrinsic cellular mechanisms underlying theta phase precession of hippocampal neurons. Trends Neurosci 30:325–333.

McKenzie S, Huszár R, English DF, Kim K, Christensen F, Yoon E, Buzsáki G (2021) Preexisting hippocampal network dynamics constrain optogenetically induced place fields. Neuron 109:1040–1054.e7.

Mehta MR, Lee AK, Wilson MA (2002) Role of experience and oscillations in transforming a rate code into a temporal code. Nature 417:741–746.

Mizuseki K, Buzsáki G (2013) Preconfigured, skewed distribution of firing rates in the hippocampus and entorhinal cortex. Cell Rep 4:1010–1021.

Mizuseki K, Sirota A, Pastalkova E, Buzsáki G (2009) Theta oscillations provide temporal windows for local circuit computation in the entorhinal-hippocampal loop. Neuron 64:267–280.

Noked O, Levi A, Someck S, Amber-Vitos O, Stark E (2021) Bidirectional optogenetic control of inhibitory neurons in freely-moving mice. IEEE Trans Biomed Eng 68:416–427.

O’Keefe J, Recce ML (1993) Phase relationship between hippocampal place units and the EEG theta rhythm. Hippocampus 3:317–330.

Pachitariu M, Steinmetz N, Kadir S, Carandini M, D HK (2016) Kilosort: realtime spike-sorting for extracellular electrophysiology with hundreds of channels. Available at: https://www.biorxiv.org/content/10.1101/061481v1 [Accessed October 17, 2021].

Pastalkova E, Itskov V, Amarasingham A, Buzsáki G (2008) Internally generated cell assembly sequences in the rat hippocampus. Science 321:1322–1327.

Qasim SE, Fried I, Jacobs J (2021) Phase precession in the human hippocampus and entorhinal cortex. Cell 184:3242–3255.e10.

Ravassard P, Kees A, Willers B, Ho D, Aharoni DA, Cushman J, Aghajan ZM, Mehta MR (2013) Multisensory control of hippocampal spatiotemporal selectivity. Science 340:1342–1346.

Reifenstein ET, Bin Khalid I, Kempter R (2021) Synaptic learning rules for sequence learning. eLife 10:e67171.

Robinson NTM, Priestley JB, Rueckemann JW, Garcia AD, Smeglin VA, Marino FA, Eichenbaum H (2017) Medial entorhinal cortex selectively supports temporal coding by hippocampal neurons. Neuron 94:677–688.e6.

Rossant C, Kadir SN, Goodman DFM, Schulman J, Hunter MLD, Saleem AB, Grosmark A, Belluscio M, Denfield GH, Ecker AS, Tolias AS, Solomon S, Buzski G, Carandini M, Harris KD (2016) Spike sorting for large, dense electrode arrays. Nat Neurosci 19:634–641.

Royer S, Zemelman BV, Losonczy A, Kim J, Chance F, Magee JC, Buzsáki G (2012) Control of timing, rate and bursts of hippocampal place cells by dendritic and somatic inhibition. Nat Neurosci 15:769– 775.

Schmidt R, Diba K, Leibold C, Schmitz D, Buzsáki G, Kempter R (2009) Single-trial phase precession in the hippocampus. J Neurosci 29:13232–13241.

Shimbo A, Izawa E-I, Fujisawa S (2021) Scalable representation of time in the hippocampus. Sci Adv 7:eabd7013.

Sirota A, Montgomery S, Fujisawa S, Isomura Y, Zugaro M, Buzsáki G (2008) Entrainment of neocortical neurons and gamma oscillations by the hippocampal theta rhythm. Neuron 60:683–697.

Skaggs WE, McNaughton BL, Wilson MA, Barnes CA (1996) Theta phase precession in hippocampal neuronal populations and the compression of temporal sequences. Hippocampus 6:149–172.

Sloin HE, Bikovski L, Levi A, Amber-Vitos O, Katz T, Spivak L, Someck S, Gattegno R, Sivroni S, Sjulson L, Stark E (2022a) Hybrid offspring of C57BL/6J mice exhibit improved properties for neurobehavioral research. eNeuro 9:ENEURO.0221-22.2022.

Sloin HE, Levi A, Someck S, Spivak L, Stark E (2022b) High fidelity theta phase rolling of CA1 neurons. J Neurosci 42:3184–3196.

Stark E, Eichler R, Roux L, Fujisawa S, Rotstein HG, Buzsáki G (2013) Inhibition-induced theta resonance in cortical circuits. Neuron 80:1263–1276.

Stark E, Koos T, Buzsáki G (2012) Diode probes for spatiotemporal optical control of multiple neurons in freely moving animals. J Neurophysiol 108:349–363.

Stark E, Roux L, Eichler R, Senzai Y, Royer S, Buzsáki G (2014) Pyramidal cell-interneuron interactions underlie hippocampal ripple oscillations. Neuron 83:467–480.

Terada S, Sakurai Y, Nakahara H, Fujisawa S (2017) Temporal and rate coding for discrete event sequences in the hippocampus. Neuron 94:1248–1262.

Tsodyks MV, Skaggs WE, Sejnowski TJ, McNaughton BL (1996) Population dynamics and theta rhythm phase precession of hippocampal place cell firing: a spiking neuron model. Hippocampus 6:271– 280.

Valero M, Zutshi I, Yoon E, Buzsáki G (2022) Probing subthreshold dynamics of hippocampal neurons by pulsed optogenetics. Science 375:570–574.

Van Der Meer MAA, Redish AD (2011) Theta phase precession in rat ventral striatum links place and reward information. J Neurosci 31:2843–2854.

Venditto SJC, Le B, Newman EL (2019) Place cell assemblies remain intact, despite reduced phase precession, after cholinergic disruption. Hippocampus 29:1075–1090.

Wang M, Foster DJ, Pfeiffer BE (2020) Alternating sequences of future and past behavior encoded within hippocampal theta oscillations. Science 370:247–250.

Wang Y, Romani S, Lustig B, Leonardo A, Pastalkova E (2015) Theta sequences are essential for internally generated hippocampal firing fields. Nat Neurosci 18:282–288.

Yamaguchi Y (2003) A theory of hippocampal memory based on theta phase precession. Biol Cybern 89:1– 9.

Yamaguchi Y, Aota Y, McNaughton BL, Lipa P (2002) Bimodality of theta phase precession in hippocampal place cells in freely running rats. J Neurophysiol 87:2629–2642.

Zugaro MB, Monconduit L, Buzsáki G (2005) Spike phase precession persists after transient intrahippocampal perturbation. Nat Neurosci 8:67–71.

